# Evolutionary optimization in classification of early-MCI patients from healthy controls using graph measures of resting-state fMRI

**DOI:** 10.1101/2021.03.04.433989

**Authors:** Jafar Zamani, Ali Sadr, Amir-Homayoun Javadi

## Abstract

Identifying individuals with early mild cognitive impairment (EMCI) can be an effective strategy for early diagnosis and delay the progression of Alzheimer’s disease (AD). Many approaches have been devised to discriminate those with EMCI from healthy control (HC) individuals. Selection of the most effective parameters has been one of the challenging aspects of these approaches. In this study we suggest an optimization method based on five evolutionary algorithms that can be used in optimization of neuroimaging data with a large number of parameters. Resting-state functional magnetic resonance imaging (rs-fMRI) measures, which measure functional connectivity, have been shown to be useful in prediction of cognitive decline. Analysis of functional connectivity data using graph measures is a common practice that results in a great number of parameters. Using graph measures we calculated 1155 parameters from the functional connectivity data of HC (n=36) and EMCI (n=34) extracted from the publicly available database of the Alzheimer’s disease neuroimaging initiative database (ADNI). These parameters were fed into the evolutionary algorithms to select a subset of parameters for classification of the data into two categories of EMCI and HC using a two-layer artificial neural network. All algorithms achieved classification accuracy of 94.55%, which is extremely high considering single-modality input and low number of data participants. These results highlight potential application of rs-fMRI and efficiency of such optimization methods in classification of images into HC and EMCI. This is of particular importance considering that MRI images of EMCI individuals cannot be easily identified by experts.

## 1 INTRODUCTION

Alzheimer’s disease (AD) is the most common type of dementia, with around 50 million patients worldwide ^1,2^. AD is usually preceded by a period of mild cognitive impairment (MCI) ^3,4^. Identifying the subjects with MCI could be an effective strategy for early diagnosis and delay the progression of AD towards irreversible brain damage ^5–7^. While researchers were successful, to some extent, in diagnosis of AD, researchers were significantly less successful in diagnosis of MCI ^8–11^. In particular, detection of early stages of MCI (EMCI) has been proven to be very challenging ^12–14^. Therefore, in this study we propose a novel method based on evolutionary algorithms to select a subset of graph features calculated from functional connectivity data to discriminate between healthy participants (HC) and EMCI.

It has been shown that the brain goes through many functionally and physiologically changes prior to any obvious behavioral symptoms in AD ^15–17^. Therefore, many approaches have been devised based on biomarkers to distinguish between HC, and different stages of MCI, and AD ^18–20^. For example, segmentation of structural magnetic resonance imaging (MRI) data has been used in many studies as brain structure changes greatly in AD ^21–24^.

While structural neuroimaging has shown some success in early detection of AD, functional neuroimaging has proven to be a stronger candidate ^25–27^. Functional MRI (fMRI) allows for the examination of brain functioning while a patient is performing a cognitive task. This technique is especially well suited to identifying changes in brain functioning before significant impairments can be detected on standard neuropsychological tests, and as such is sensitive to early identification of the disease processes ^28,29^. While fMRI requires participants to perform a task, resting-state fMRI (rs-fMRI) is capable of measuring the spontaneous fluctuations of brain activity without any task, hence it is less sensitive to individual cognitive abilities ^30–32^. One important feature of rs-fMRI is the ability to measure functional connectivity changes ^33,34^ with many recent studies have shown that functional connectivity changes are prevalent in AD ^35–38^.

Analysis of rs-fMRI data using graph theory measures is a powerful tool that enables characterization of the global, as well as local, characteristics of different brain areas ^39–42^. This method provides us with a way to comprehensively compare functional connectivity organization of the brain between patients and controls ^43–45^, and has been used in characterization of AD ^46–48^. This method has also been used in diagnosis classification of different stages of AD ^49–51^.

Since graph theory analysis of rs-fMRI data leads to a large number of parameters, it is essential to select an optimal subset of features that can lead to high discrimination accuracy ^52,53^. Feature selection is particularly complicated due to the non-linear nature of classification methods. For example, more parameters do not necessarily lead to better performance ^54,55^. Evolutionary algorithms (EA) are biologically-inspired algorithms that are extremely effective in optimization algorithms with large search spaces ^56–60^. EA has been used in characterization and diagnosis of AD ^61–65^.

In this study we devised a method that achieves higher accuracy in the classification of HC and EMCI participants compared to the past published research. We used MRI and rs-fMRI data of a group of healthy participants and those with EMCI. We applied graph theory to extract a collection of 1155 parameters. This data is then given to five different EA methods to select an optimum subset of parameters. These selected parameters are subsequently given to an artificial neural network to classify the data into two groups of HC and EMCI. We aimed at identifying the most suitable method of optimization based on accuracy and training time, as well as identifying the most informative parameters.

## 2 Methods

### 2.1 Participants

Data for 70 participants were extracted from the publicly available database of the Alzheimer’s disease neuroimaging initiative database (ADNI) (http://adni.loni.usc.edu) ^66–68^. Table 1 represents the details of the data. EMCI participants had no other neurodegenerative diseases except MCI. The EMCI participants were recruited with memory function approximately 1.0 SD below expected education adjusted norms ^69^. HC subjects had no history of cognitive impairment, head injury, major psychiatric disease, or stroke.

**Table 1.** Demographics of the data for participants included in this study.

|  | **EMCI** | **HC** | ***p*** |
| --- | --- | --- | --- |
| n | 34 | 36 |  |
| Female (n [%]) | 19 [55.88] | 19 [52.77] |  |
| Age (mean [SD]) | 71.73 [7.80] | 69.97 [5.60] | 0.297 |
| MMSE (mean [SD]) | 28.61 [1.60] | 28.40 [4.60] | 0.302 |
| CDR | 0.5 or 1 | 0 | < 0.001 |

notes: CDR: clinical dementia rating, MMSE: mini-mental state exam, HC: healthy control, EMCI: early mild cognitive impairment

### 2.2 Proposed Method

Structural T1-MRI and rs-fMRI data was extracted from the ADNI database ^68^. The data is given to CONN toolbox ^70^ in MATLAB v2018 (MathWorks, California, US). CONN is a tool for preprocessing, processing, and analysis of functional connectivity data. Preprocessing consisted of reducing subject motion, image distortions, and magnetic field inhomogeneity effects and application of denoising methods for reduction of physiological effects and other sources of noise. The processing stage consisted of extraction of functional connectivity and graph theory measures. In this stage, through two pipelines, a collection of 1155 parameters are extracted ^70,71^. These parameters are then given to one of the dimension reduction methods (five EA and one statistical method) to select a subset of features. The selected features are finally given to an artificial neural network to classify the data into two categories of healthy control (HC) and EMCI. See Figure 1 for the summary of the procedure of the method.

**Figure 1.**
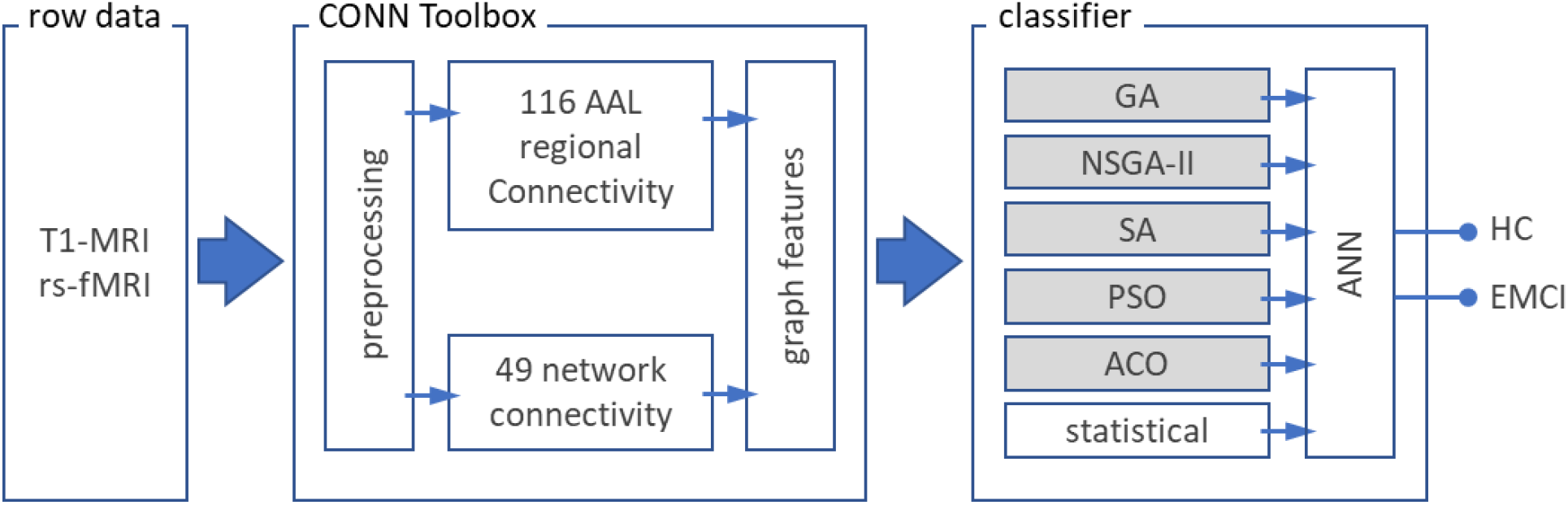
Procedure of the proposed method. T1-MRI and resting-state fMRI (rs-fMRI) data of healthy participants (HC; n=36) and patients with early mild cognitive impairment (EMCI; n=34) are extracted from ADNI database ^68^. Preprocessing, parcellation of brain area (116 regions based on AAL) and extraction of the functional connectivity (49 network parameter), as well as the seven graph parameters are done using CONN toolbox ^70^. Subsequently the 1155 (116×7 + 49×7) extracted parameters are given to one of the optimization methods to select the best subset of parameters that lead to best classification method. Optimization methods consisted of five evolutionary algorithms (boxes with grey shading) and one statistical algorithm. The outputs of these methods are given to an artificial neural network (ANN) with two hidden layers to classify the data into HC and EMCI. AAL: automated anatomical atlas ^75^; GA: genetic algorithm; NSGA-II: nondominated sorting genetic algorithm II; ACO: ant colony optimization; SA: simulated annealing; PSO: particle swarm optimization; seven graph features: degree centrality, betweenness centrality, path length, clustering coefficient, local efficiency, cost and global efficiency.

### 2.3 Data acquisition and preprocessing

Brain structural T1-weighted MRI data with 256×256×170 voxels and 1×1×1 mm^3^ voxel size were extracted for all subjects. MRI data preprocessing steps consisted of non-uniformity correction, segmentation into grey matter, white matter and cerebrospinal fluid (CSF) and spatial normalization to MNI space.

Using an echo-planar imaging sequence on a 3T Philips MRI scanner, rs-fMRI data were obtained. Acquisition parameters were: 140 time points, repetition time (TR) = 3000 ms, echo time (TE) = 30 ms, flip angle = 80°, number of slices = 48, slice thickness= 3.3 mm, spatial resolution = 3×3×3 mm^3^ and in plane matrix = 64×64. fMRI images preprocessing steps consisted of motion correction, slice timing correction, spatial normalization to MNI space, low frequency filtering to keep only (0.01 – 0.1 Hz) fluctuations. T1-MRI and rs-fMRI data processing was done using CONN toolbox ^70^.

### 2.4 Functional Connectivity

Functional connectivity, also called “resting state” connectivity, is a measure for the temporal correlations among the blood-oxygen-level-dependent (BOLD) signal fluctuations in different brain areas ^72–74^. In this study, we obtained functional connectivity of region of interest (ROI)-to-ROI of brain areas according to Harvard-Oxford atlas ^75^. The functional connectivity matrix is the correlation, covariance, or the mutual information between the fMRI time series of every two brain regions, which is stored in an *n* × *n* matrix for each participant, where n is the number of brain regions obtained by atlas parcellation ^74^. To extract functional connectivity between different brain areas we used Pearson correlation coefficients formula as following ^70,76^:

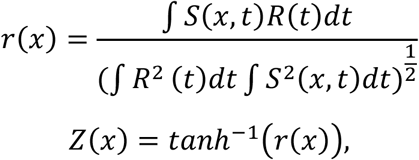

where *S* is the BOLD time series at each voxel (for simplicity all-time series are considered central to zero means), *R* is the average BOLD time series within an ROI, *r* is the spatial map of Pearson correlation coefficients, and *Z* is the seed-based correlations (SBC) map of Fisher-transformed correlation coefficients for this ROI ^77^.

### 2.5 Graph Parameters

We used the graph theory technique to study topological features of functional connectivity graphs across multiple regions of the brain ^49,78^. Graph nodes represented brain regions and edges represented interregional resting-state functional connectivity. The functional connectivity matrix is employed for estimating common features of graphs including (1) *degree centrality* (the number of edges that connect a node to the rest of the network) (2) *betweenness centrality* (the proportion of shortest paths between all node pairs in the network that pass through a given index node), (3) *average path length* (the average distance from each node to any other node), (4) *clustering coefficient* (the proportion of ROIs that have connectivity with a particular ROI that also have connectivity with each other), (5) *cost* (the ratio of the existing number of edges to the number of all possible edges in the network), (6) *local efficiency* (the network ability in transmitting information at the local level), (7) *global efficiency* (the average inverse shortest path length in the network; this parameter is inversely related to the path length) ^79^.

### 2.6 Dimension Reduction Methods

We used five EA to select the most efficient set number of features. These algorithms are as follows:

#### Genetic algorithm (GA)

GA is one of the most advanced algorithms for feature selection ^80^. This algorithm is based on the mechanics of natural genetics and biological evolution for finding the optimum solution. It consists of five steps: selection of initial population, evaluation of fitness function, pseudo-random selection, crossover, and mutation ^81^. For further information refer to supplementary Methods section. Single point, double point, and uniform crossover methods are used to generate new members. In this study we used 0.3 and 0.1 as mutation percentage and mutation rate, respectively; 20 members per population, crossover percentage was 14 with 8 as selection pressure ^63,82^.

#### Nondominated sorting genetic algorithm II (NSGA-II)

NSGA is a method to solve multi-objective optimization problems to capture a number of solutions simultaneously ^83^. All the operators in GA are also used here. NSGA-II uses binary features to fill a mating poll. Nondomination and crowding distance are used to sort the new members. For further information refer to supplementary Methods section. In this study the mutation percentage and mutation rate were set to 0.4 and 0.1, respectively; population size was 25, and crossover percentage was 14%.

#### Ant colony optimization algorithm (ACO)

ACO is a metaheuristic optimization method based on the behavior of ants ^84^. This algorithm consists of four steps: initialization, creation of ant solutions (a set of ants build a solution to the problem being solved using pheromones values and other information), local search (improvement of the created solution by ants), and global pheromone update (update in pheromone variables based on search action followed by ants) ^85^. ACO requires a problem to be described as a graph: nodes represent features and edges indicate which features should be selected for the next generation. In features selection, the ACO tries to find the best solutions using prior information from previous iterations. The search for the optimal feature subset consists of an ant traveling through the graph with a minimum number of nodes required for satisfaction of stopping criterion ^86^. For further information refer to supplementary Methods section. We used 10, 0.05, 1, 1 and 1 for the number of ants, evaporation rate, initial weight, exponential weight, and heuristic weight, respectively.

#### Simulated annealing (SA)

SA is a stochastic search algorithm, which is particularly useful in large-scale linear regression models ^87^. In this algorithm, the new feature subset is selected entirely at random based on the current state. After an adequate number of iterations, a dataset can be created to quantify the difference in performance with and without each predictor ^88,89^. For further information refer to supplementary Methods section. We set initial temperature and temperature reduction rate with 10 and 0.99, respectively.

#### Particle swarm optimization (PSO)

PSO is a stochastic optimization method based on the behavior of swarming animals such as birds and fish. Each member finds optimal regions of the search space by coordinating with other members in the population. In this method, each possible solution is represented as a particle with a certain position and velocity moving through the search space ^90–92^. Particles move based on cognitive parameter (defining the degree of acceleration towards the particle’s individual local best position, and global parameter (defining the acceleration towards the global best position). The overall rate of change is defined by an inertia parameter. For further information refer to supplementary Methods section. We used 20 as the warm size, cognitive and social parameters were set to 1.5 and inertia as 0.72.

#### Statistical approach

To create a baseline to compare dimension reduction methods based on evolutionary algorithms, we also used the statistical approach to select the features based on the statistical difference between the two groups. We compared the 1155 parameters using two independent-sample t-test analyses. Subsequently we selected the parameters based on their sorted *p* values.

### 2.7 Classification Method

For classification of EMCI and HC we used a multi-layer perceptron artificial neural network (ANN) with two fully-connected hidden layers with 10 nodes each. Classification method was performed via a 10-fold cross-validation. We used Levenberg-Marquardt Back propagation (LMBP) algorithm for training ^93–95^ and mean square error as a measure of performance. The LMBP has three steps: (1) propagate the input forward through the network; (2) propagate the sensitivities backward through the network from the last layer to the first layer; and finally (3) update the weights and biases using Newton’s computational method ^93^. In the LMBP algorithm the performance index *F*(*x*) is formulated as:

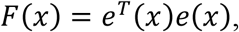

where *e* is vector of network error, and *x* is the vector matrix of network weights and biases. The network weights are updated using the Hessian matrix and its gradient:

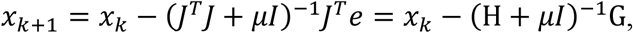

where *J* represent Jacobian matrix. The Hessian matrix *H* and its gradient *G* are calculated using:

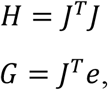

where the Jacobian matrix is calculated by:

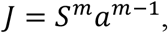

where *a*^*m*−1^ is the output of the (*m* − 1)th layer of the network, and *S*^*m*^ is the sensitivity of *F*(*x*) to changes in the network input element in the *m*th layer and is calculated by:

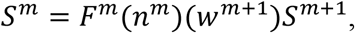

where *w*^*m*+1^ represents the neuron weight at (*m* + 1)th layer, and *n* is the network input ^93^.

## 3 RESULTS

The preprocessing and processing of the data was successful. We extracted 1155 graph parameters per participant (see Supplementary Figures 1-11). This data was used for the data optimization step. Using the five EA optimization methods and the statistical method, we investigated the performance of the classification for different numbers of subset of parameters. Figure 2 shows the performance of these methods for 100 subsets of parameters with 1 to 100 parameters. These plots are created based on 200 repetitions of the EA algorithms. To investigate the performance of the algorithms with more repetitions, we ran the same algorithms with 500 repetitions. These simulations showed no major improvement of increased repetition (maximum 0.84% improvement; see Supplementary Figure 12).

**Figure 2.**
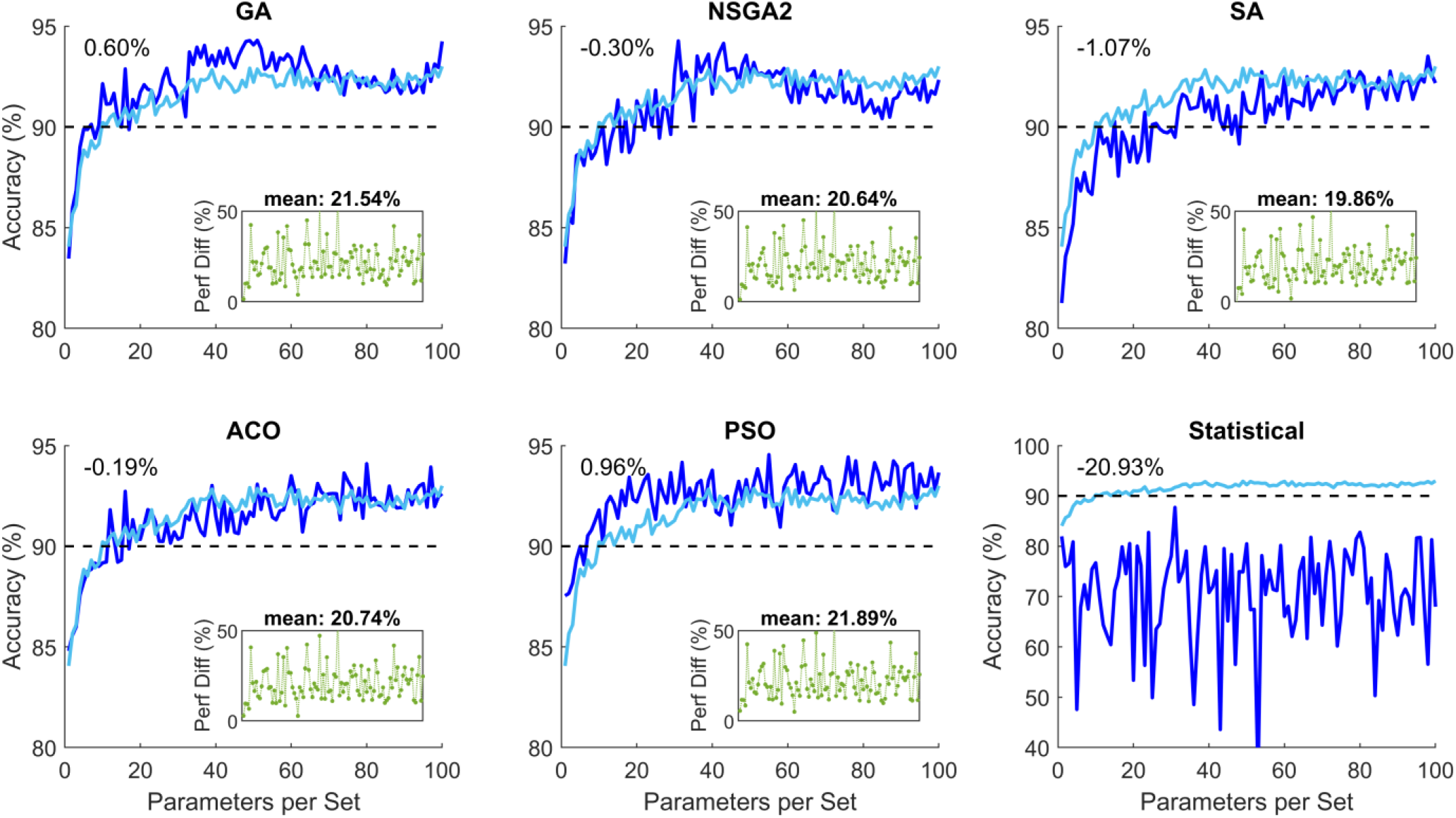
Classification performance of the five evolutionary algorithm (EA) methods and the statistical method for parameter subsets with 1 to 100 elements. The light blue color shows the average of the five EV algorithms. The number on the top left-hand corner represents the difference between the relevant plot and the mean performance of the EA methods. The green plot subplot in each panel represents superiority of the relevant EA as compared to the statistical method for different 100 subsets. The percentage value above the subplot shows the mean superior performance for the 100 subsets compared to the statistical method. These plots show that the EA performed significantly better than the statistical method. GA: genetic algorithm; NSGA-II: nondominated sorting genetic algorithm II; ACO: ant colony optimization; SA: simulated annealing; PSO: particle swarm optimization.

A threshold of 90% was chosen as the desired performance accuracy. Statistical modeling performance was constantly less than this threshold. The five EA methods achieved this performance with varying number of parameters. Figure 3 shows the accuracy percentage and the optimization speed of the five EA methods.

**Figure 3.**
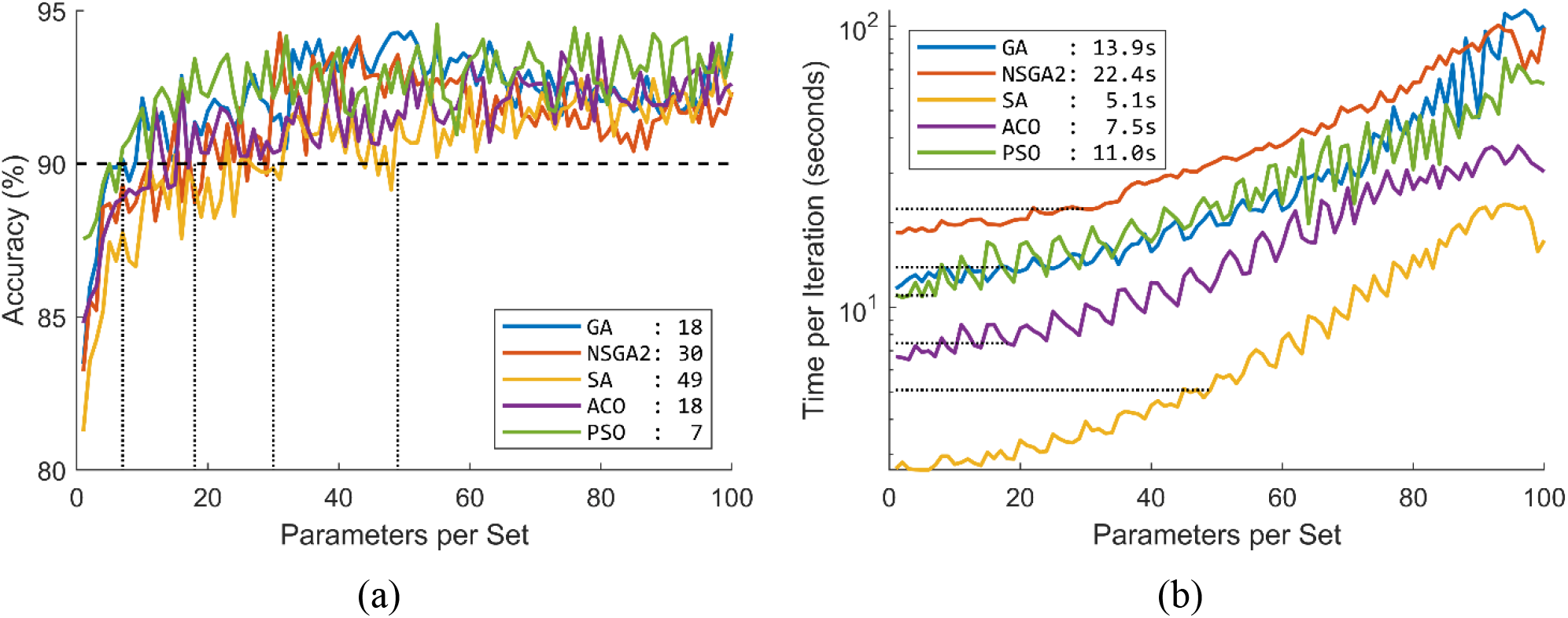
Performance of the five evolutionary algorithms (EA) in terms of (a) percentage accuracy and (b) optimization speed. The values in the legend of panel (a) show the minimum number of parameters required to achieve minimum 90% accuracy. The values in the legend of panel (b) show the minimum optimization speed to achieve minimum 90% accuracy. GA: genetic algorithm; NSGA-II: nondominated sorting genetic algorithm II; ACO: ant colony optimization; SA: simulated annealing; PSO: particle swarm optimization.

To investigate whether increasing number of parameters would increase performance, we performed similar simulations with maximum 500 parameters in each subset. This analysis showed that the performance of the optimization methods plateaus without significant increase from 100 parameters (Figure 4). This figure shows that performance of the optimization methods was between 92.55-93.35% and 94.27-94.55% for filtered and absolute accuracy, respectively. These accuracy percentages are significantly higher than 81.97% and 87.72% for filtered and absolute accuracy in the statistical classification condition.

**Figure 4.**
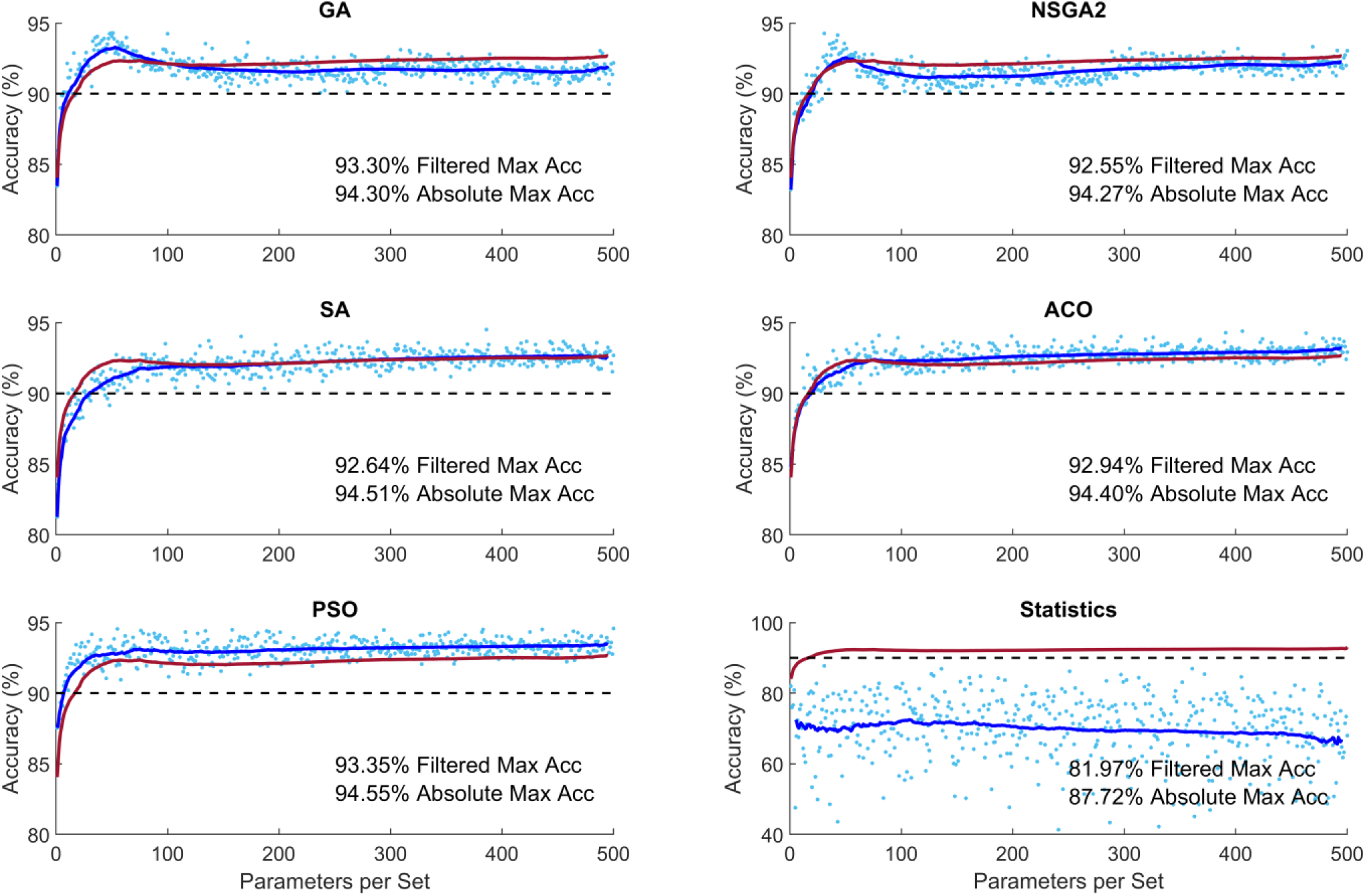
Performance of different optimization methods for increased number of parameters per subset. The light blue dots indicate the performance of algorithms for each subset of parameters. The dark blue curve shows the moving average of the samples with window of ±20 points (Filtered Data). The red curve shows the mean performance of the five evolutionary algorithms. GA: genetic algorithm; NSGA-II: nondominated sorting genetic algorithm II; ACO: ant colony optimization; SA: simulated annealing; PSO: particle swarm optimization.

To investigate the contribution of different parameters in the optimization of classification we looked at the distribution of parameters in the 100 subsets calculated above (Figure 5). GA and NSGA showed that the majority of the subsets consisted of repeated parameters: out of the 1155 parameters only about 200 of the parameters were selected in the 100 subsets. SA, ACO and PSO, on the other hand, showed a more diverse selection of parameters: almost all the parameters appeared in at least one of the 100 subsets.

**Figure 5.**
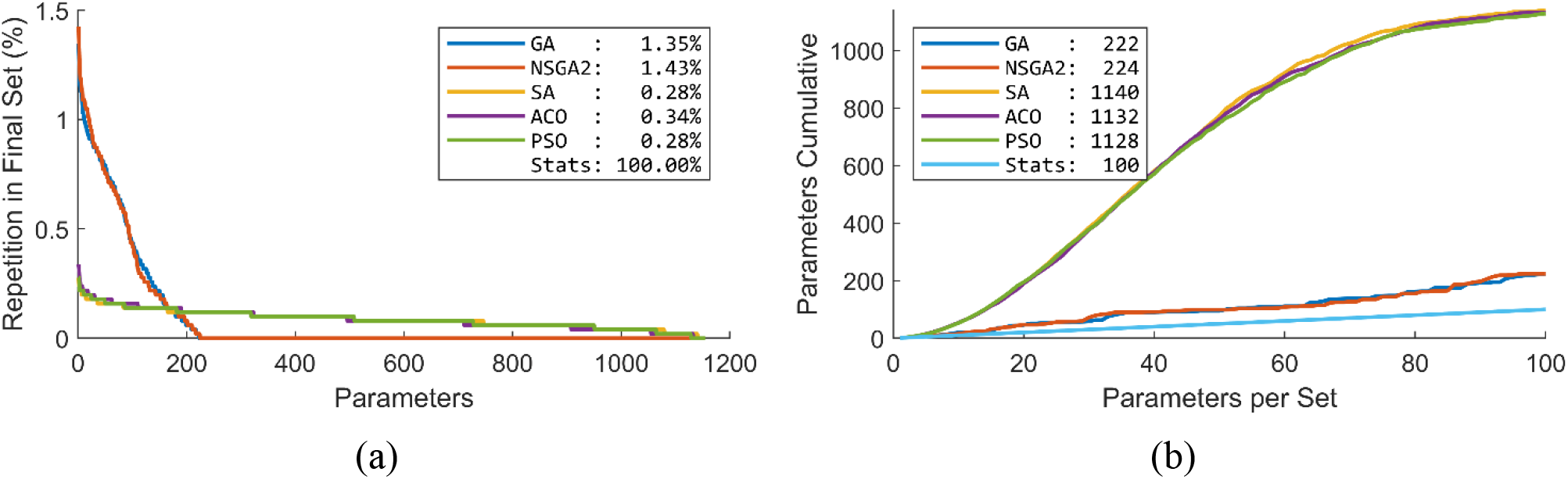
Distribution of different parameters over the 100 subsets of parameters. (a) Percentage of presence of the 1155 parameters. In the Statistical method, which is not present in the plot, the first parameter was repeated in all the 100 subsets. Numbers in the legend show the percentage repetition of the most repeated parameter. (b) Cumulative number of unique parameters over the 100 subsets of parameters. This plot shows that GA and NSGA2 concentrated on a small number of parameters, while the SA, ACO and PSO selected a more diverse range of parameters in the optimization. Numbers in the legend show the number of utilized parameters in the final solution of the 100 subsets of parameters. GA: genetic algorithm; NSGA-II: nondominated sorting genetic algorithm II; ACO: ant colony optimization; SA: simulated annealing; PSO: particle swarm optimization.

## 4 Discussions

Using CONN toolbox we extracted 1155 graph parameters from rs-fMRI data. The optimization methods showed superior performance over statistical analysis (average 20.93% superiority). The performance of the EA algorithms did not differ greatly (range 92.55-93.35% and 94.27-94.55% for filtered and absolute accuracy, respectively) with PSO performing the best (mean 0.96% superior performance) and SA performing the worst (mean 1.07% inferior performance), (Figure 2). The minimum number of required parameters to guarantee at least 90% accuracy differed quite greatly across the methods (PSO and SA requiring 7 and 49 parameters, respectively). The processing time to achieve at least 90% accuracy also differed across the EA methods (SA and NSGA2 taking 5.1s and 22.4s per optimization), (Figure 3). Increased number of parameters per subset did not increase the performance accuracy of the methods greatly, (Figure 4).

Classification of data into AD and HC has been investigated extensively. Many methods have been developed using different modalities of biomarkers. Some of these studies achieved accuracies greater than 90% ^96^. Classification of earlier stages of AD, however, has been more challenging; only a handful of studies have achieved accuracy higher than 90% (Table 2). The majority of these studies implemented convolutional and deep neural networks that require extended training and testing durations with many input data. For example, Payan et al. (2015) applied convolutional neural networks (CNN) on a collection of 755 HC and 755 MCI and achieved accuracy of 92.1% ^97^. Similarly, Wang et al. (2019) applied deep neural networks to 209 HC and 384 MCI data and achieved accuracy of 98.4% ^98^ (see also ^99–102^). We applied our method to a group of only 70 participants and achieved an accuracy of 94.55%. To the best of our knowledge, between all the studies published to date, this accuracy level is the second highest accuracy after Wang et al (2019) ^98^ with 593 total number of participants.

**Table 2.**
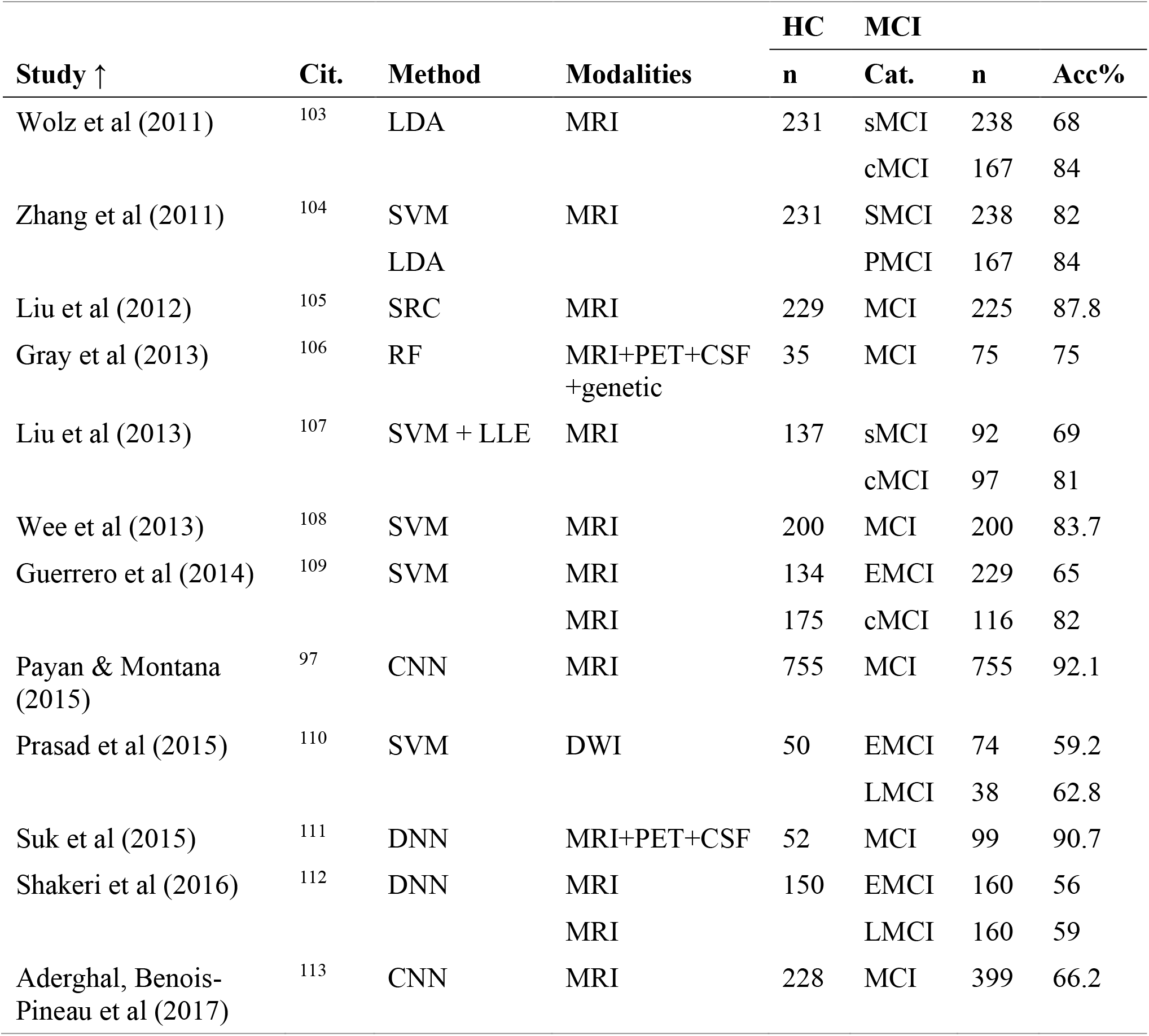
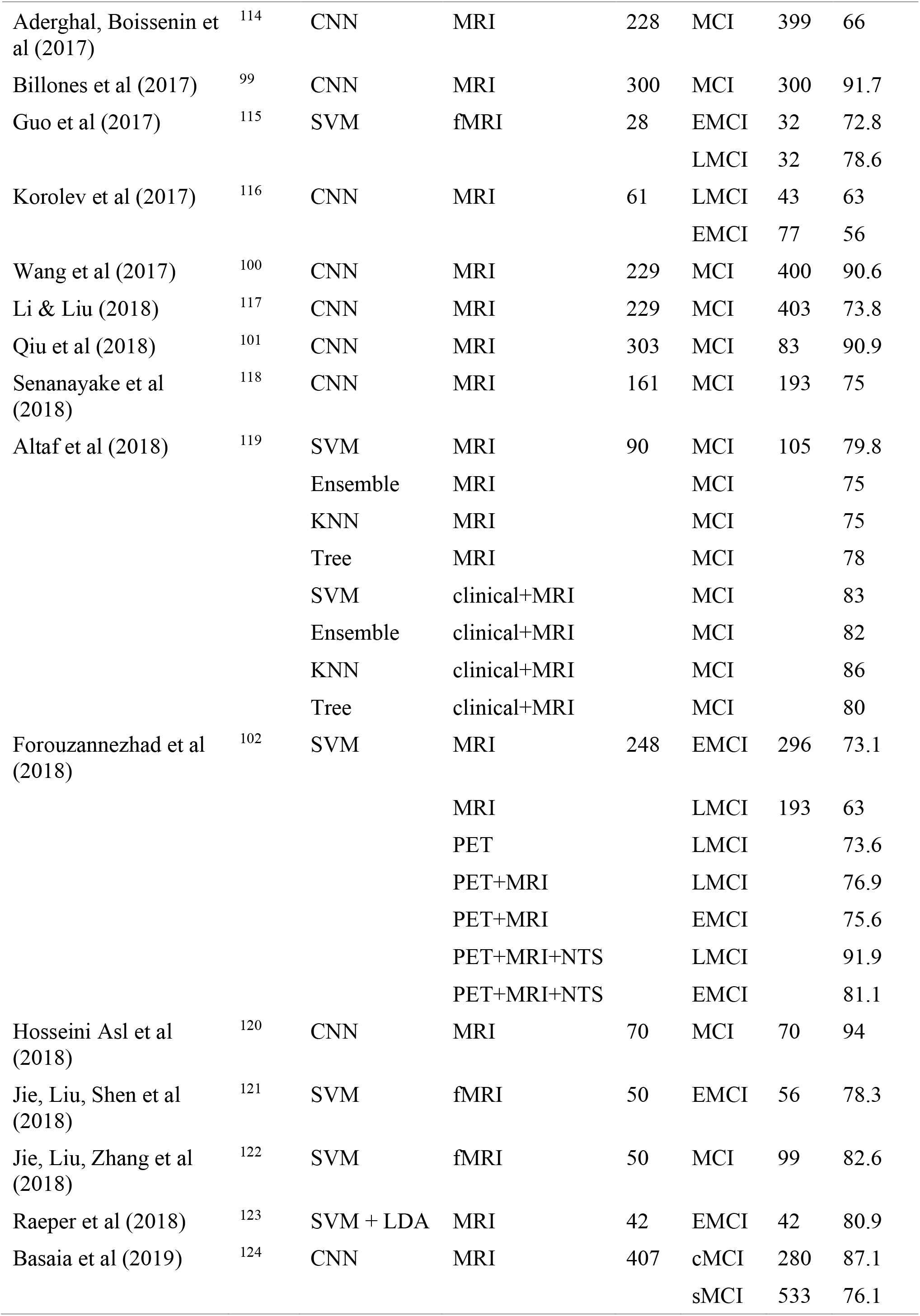
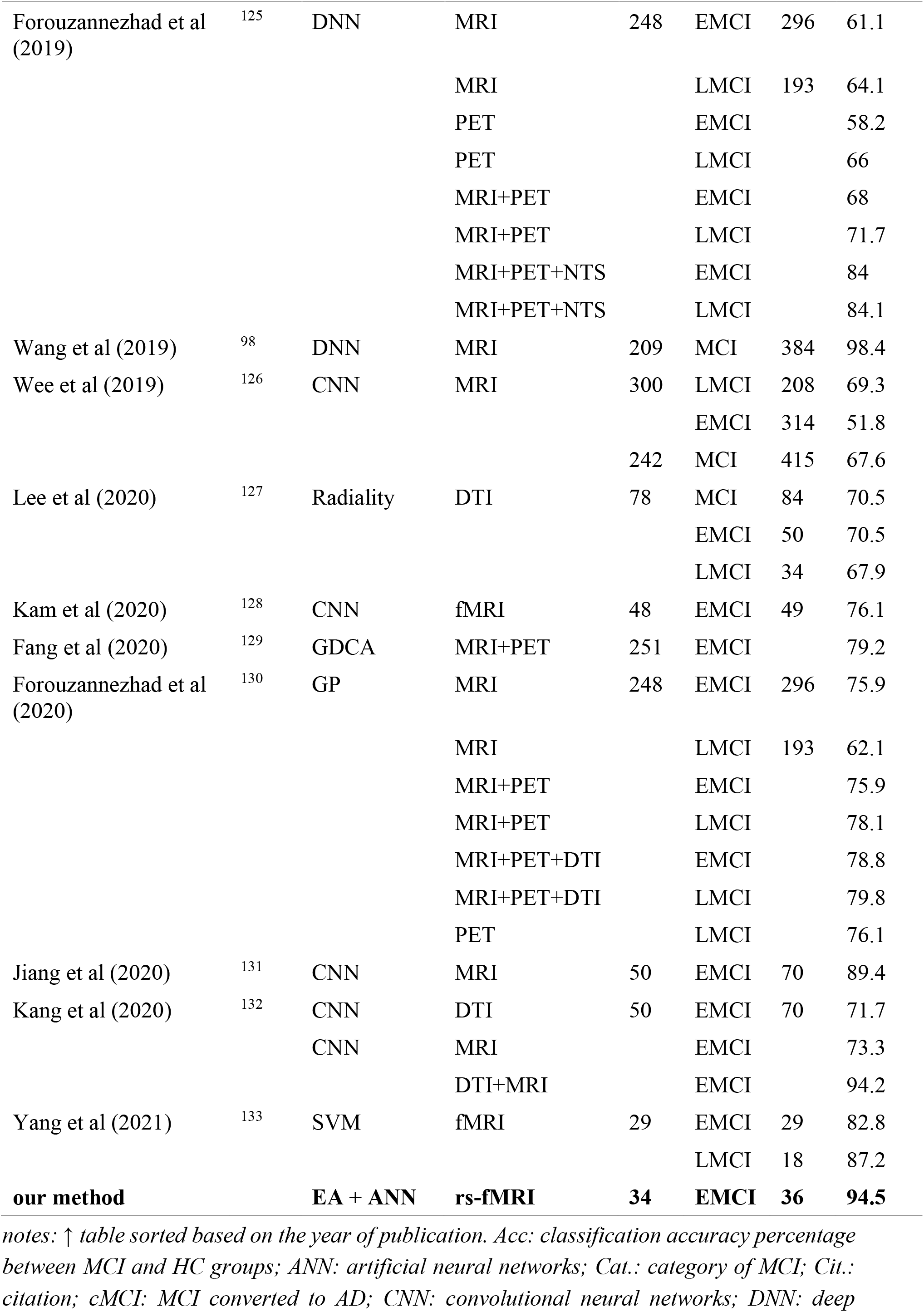
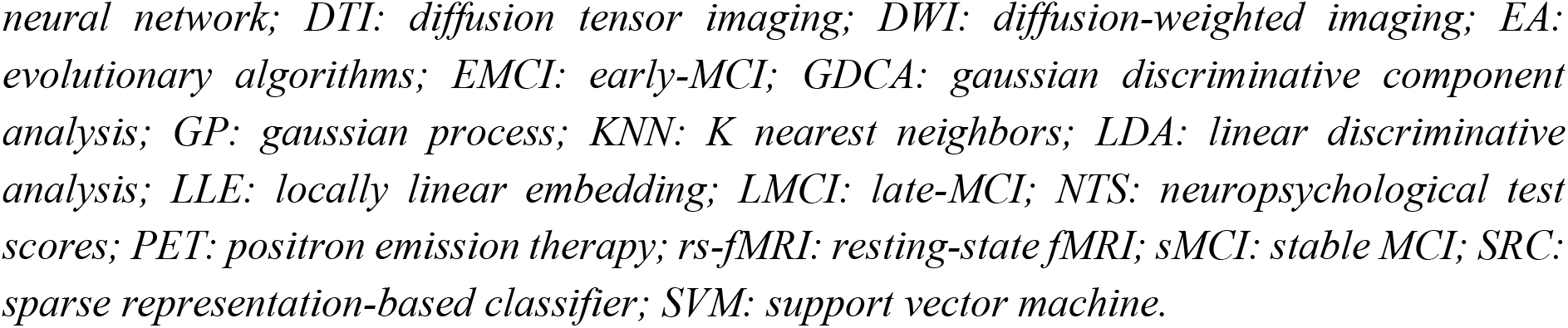
Summary of the studies aiming at categorization of healthy (HC) and mild cognitive impairment (MCI) using different biomarkers and classification methods. Only best performance of each study is reported for each group of participants and classification method. Further details of the following studies are in Supplementary Table 1.

Research has shown that having a combination of information from different modalities supports higher classification accuracies. For example, Forouzannezhad et al. (2018) showed that a combination of PET, MRI and neuropsychological test scores (NTS) can improve performance by more than 20% as compared to only PET or MRI ^102^. In another study, Kang et al. (2020) showed that a combination of diffusion tensor imaging (DTI) and MRI can improve accuracy by more than 20% as compared to DTI and MRI alone ^132^. Our analysis, while achieving superior accuracy compared to a majority of the prior methods, was based on one biomarker of MRI, which has a lower computational complexity than multi-modality data.

Interpretability of the selected features is one advantage of the application of evolutionary algorithms as the basis of the optimization algorithm. This is in contrast with algorithms based on CNN or deep neural networks (DNN) that are mostly considered as black boxes ^134^. Although research has shown some progress in better understanding the link between the features used by the system and the prediction itself in CNN and DNN, such methods remain difficult to verify ^135,136^. This has reduced trust in the internal functionality and reliability of such systems in clinical settings ^137^. Our suggested method clearly selects features based on activity of distinct brain areas, which are easy to interpret and understand ^64,138^. This can inform future research by bringing the focus to brain areas and the link between brain areas that are more affected by mild cognitive impairment.

We implemented five of the most common evolutionary algorithms. They showed similar overall optimization performance ranging between 92.55-93.35% and 94.27-94.55% for filtered and absolute accuracy, respectively. They, however, differed in optimization curve, optimization time and diversity of the selected features. PSO could guarantee a 90% accuracy with only 7 features. SA on the other hand required 49 features to guarantee a 90% accuracy. Although SA required more features to guarantee a 90% accuracy, it was the fastest optimization algorithm with only 5.1s for 49 features. NSGA2 on the other hand required 22.4s to guarantee a 90% accuracy. These show the diversity of the algorithms and their suitability in different applications requiring highest accuracy, least number of features or fastest optimization time ^56,61,139^.

One distinct characteristic of GA and NSGA-II was the more focused search amongst features as compared to the other methods. GA and NSGA-II selected 222 and 224 distinct features in the first 100 parameter sets, respectively, while the other methods covered almost the whole collection of features, covering more than 97.6%. Notably GA and NSGA-II showed “curse of dimensionality” (also known as “peaking phenomenon”) with optimal number of features around 50 parameters ^140–143^. Therefore, perhaps the features selected by GA and NSGA-II are more indicative of distinct characteristics of the differences between HC and EMCI.

In this study, we proposed a method for classification of the EMCI and HC groups using graph theory. These results highlight the potential application of graph analysis of functional connectivity and efficiency of evolutionary algorithm in combination with a simple perceptron ANN in the classification of images into HC and EMCI. We proposed a fully automatic procedure for predication of early stages of AD using rs-fMRI data features. This is of particular importance considering that MRI images of EMCI individuals cannot be easily identified by experts. Further development of such methods can prove to be a powerful tool in the early diagnosis of AD.

## Acknowledgements

The authors would like to thank Oliver Herdson for his comments and proofreading the manuscript.

## Authors Contribution

JZ and AHJ conceived the study. JZ extracted the data. JZ and AHJ analyzed the data. JZ and AHJ wrote the paper. JZ, AS and AHJ revised the manuscript. AS and AHJ supervised the project.

## Supplementary Methods

### 1 Genetic algorithm (GA)

The procedure of GA consists of the following four steps ^1^:

1. *Individual encoding*: Each individual is encoded as binary vector of size *P*, where the entry *b*_*i*_ = 1 states for the predictor *p*_*i*_ that is defined for that individual, *b*_*i*_ = 0 if the predictor *p*_*i*_ is not included in that particular individual (*i* = 1,…, *P*).
2. *Initial population*: Given the binary representation of the individuals, the population is a binary matrix where its rows are the randomly selected individuals, and the columns are the available predictors. An initial population with a predefined number of individuals is generated with a random selection of 0 and 1 for each entry.
3. *Fitness function*: the fitness value of the individual in the population is calculated using predefined fitness function. Individual with the lowest prediction error and fewer predictors have been selected for next generation.
4. *Genetic operators*: applying genetic operators to create the next generation.

The genetic operators are, *Selection* (randomly selection of members based on their fitness value; fitter members are more likely to be chosen), *Crossover* (the new generation is created by exchanging elements between two selected parents from the previous step), *Mutation* (elements in a selected member is changed), and *Stop Criteria* (the criteria and indicate the end of the search) ^1^. In our study we used roulette wheel selection for selection of the possible valuable solutions to producing offsprings for the next generation.

### 2 Nondominated sorting genetic algorithm II (NSGA-II)

Nondomination and crowding distance are used to sort the new members. A specific number of individuals in the sorted population are transferred to the next generation. This conventional NSGA algorithm has a computational complexity of *O*(*MN*^3^), where *M* is the number of objectives and *N* is the population size. NSGA-II on the other hand has overall complexity *O*(*MN*^2^), which is significantly ^2^. After termination of the optimization process, nondominated solutions form the Pareto frontier. Each of the solutions on the Pareto frontier can be considered as an optimal strategy for a specific situation ^3–5^.

### 3 Ant colony optimization algorithm (ACO)

See Figure 1 for the procedure of the traverse of an ant placed at node *a*. This ant has a choice of which feature to add next to its path (dotted lines). It traverses through the graph to find a path that satisfies the stopping criterion (e.g., a suitably high classification accuracy has been achieved with this subset). In this example, the ant chooses next feature *b* based on a set of transition rules, then *c* and then *d*. Upon arrival at *d*, the current subset {*a*; *b*; *c*; *d*} is determined to satisfy the traversal stopping criterion. At termination of search, the algorithm outputs this feature subset as a candidate for data reduction ^6^.

**Figure 1.**
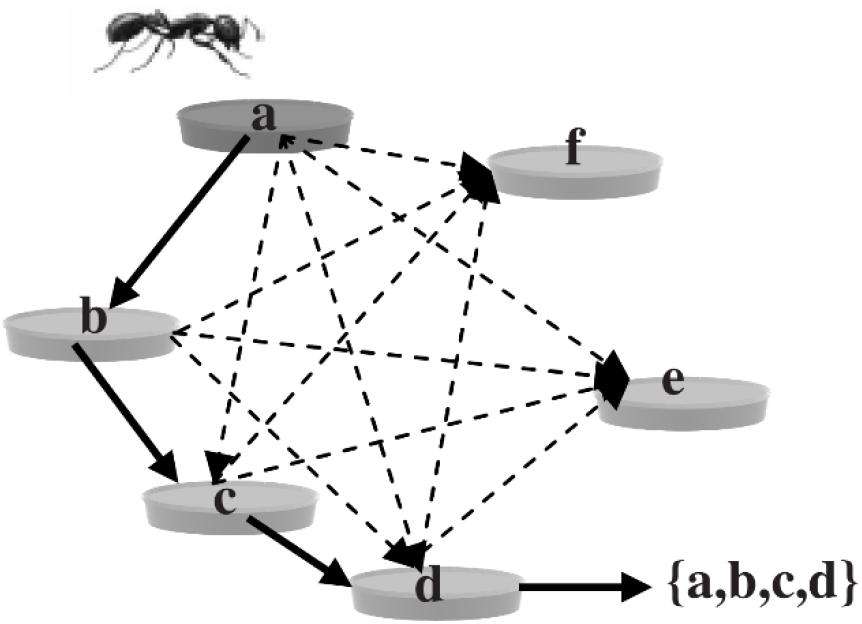
A sample example of ant traveling through multiple features in ant colony optimization algorithm (ACO). Here feature subset of {*a*; *b*; *c*; *d*} is selected as a possible solution ^6^.

The probability of an ant at feature *i* choosing to travel to feature *j* at time *t*:

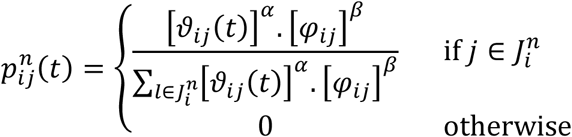

where *n* is the number of ants, *φ*_*ij*_ is the heuristic desirability of choosing feature *j* when at feature *i*, 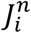 is the set of nodes next to node *i*, which have not yet been visited by the ant *n*. The *α* > 0 and *β* > 0 are two parameters that determine the relative importance of the pheromone value and heuristic information, respectively, and *ϑ*_*ij*_ is the amount of virtual pheromone on edge (*i*, *j*). The pheromone on each edge is updated according to the following formula ^6^:

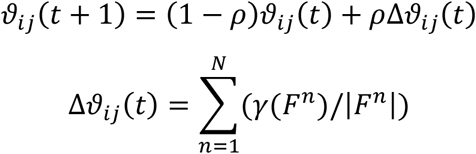

This is the case if the edge (*i*, *j*) has been traversed; Δ*ϑ*_*ij*_(*t*) is 0 otherwise. The value 0 ≤ *ρ* ≤ 1 is decay constant used to simulate the evaporation of the pheromone. The pheromone is updated according to both the measure of the “goodness” of the ant’s feature subset *γ* and the size of the subset itself. By this definition, all ants update the pheromone ^6^. *F*^*n*^ is the feature subset found by ant *n*.

### 4 Simulated annealing (SA)

SA utilizes a certain probability to accept a worse solution. The algorithm starts with a randomly generated solution; in each iteration, a neighbor solution to the best solution so far is generated according to a predefined neighborhood structure and evaluated using a fitness function. The improving move is accepted, whilst worse neighbors are accepted with a certain probability determined by the Boltzmann probability, *P* = *e* − *θ/ T* where *θ* is the difference between the fitness of the best solution and the generated neighbor. Moreover, *T* is a temperature, which periodically decreases during the search process according to a certain cooling schedule. First, the current temperature *T* is set to be a very large number ^7,8^.

### 5 Particle swarm optimization (PSO)

In a PSO with an N-dimensional search space, the particle position and velocity are formulated by:

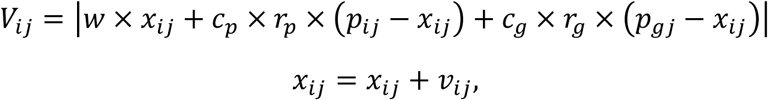

where *V*_*i*_ and *x*_*i*_ refer to the velocity and position of the particle *i*, respectively, and *j*, ranging from 1 to N (total number of features). *c*_*p*_ is the cognitive parameter, defining the degree of acceleration towards the particle’s individual local best position *p*_*ij*_. *c*_*g*_ is a social parameter, defining the acceleration towards the global best position *p*_*gj*_. *w* is an inertia parameter, regulating the overall rate of change. The stochastic nature of the velocity equation is represented by *r*_*p*_ and *r*_*g*_, which are numbers in the range [0, 1]. To maintain coherence in the swarm, the maximum velocity is regulated by a parameter *v*_*max*_. In standard PSO implementations, typically *v*_*max*_ = |*x*_*max*_ − *x*_*min*_|.

---

**Figure 1.**
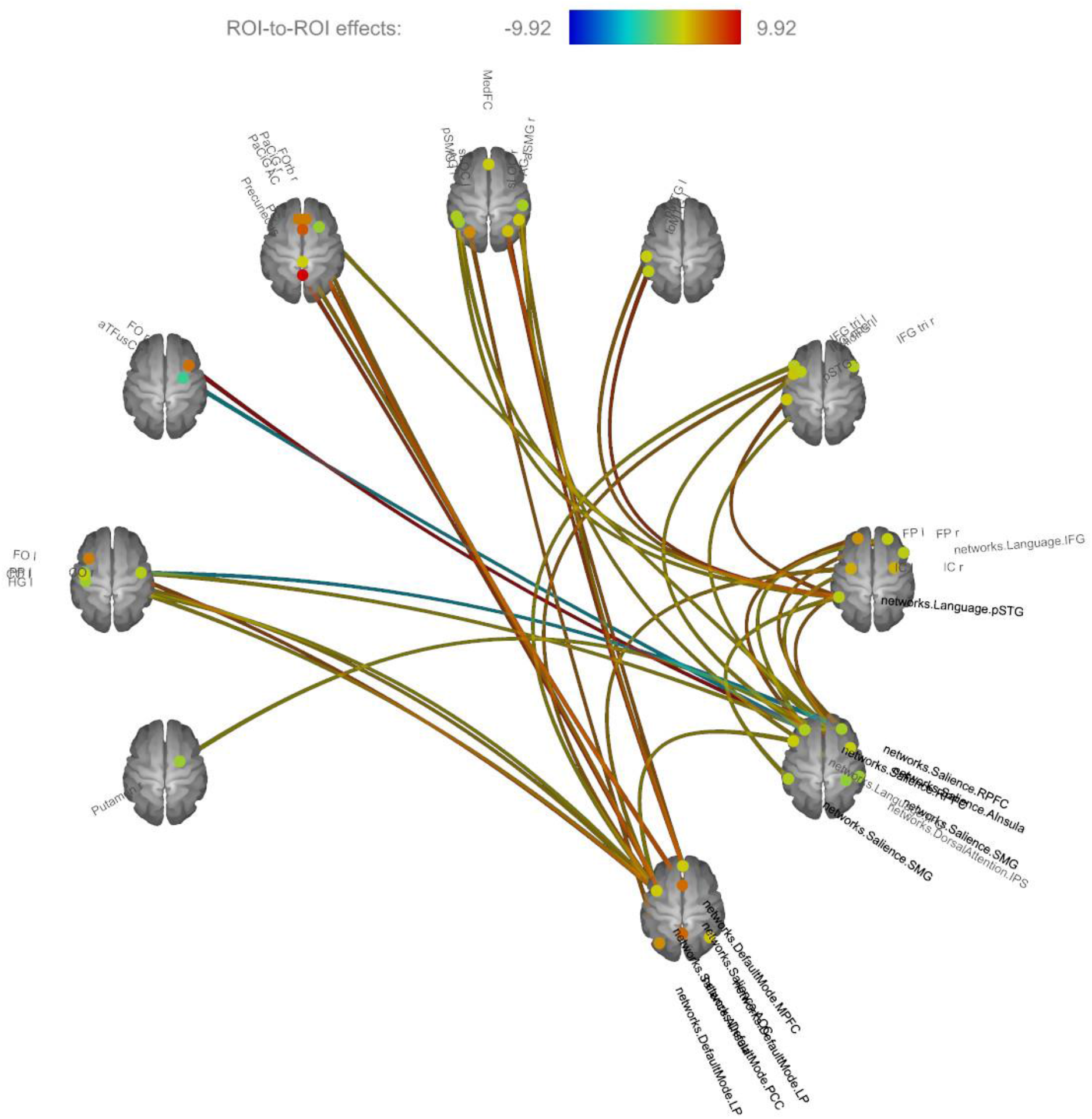
A sample collection of networks and regions of interests (ROI) connectivity matrix using rs-fMRI data. The colors indicate t-value for one-sample t-test statistics.

**Figure 2.**
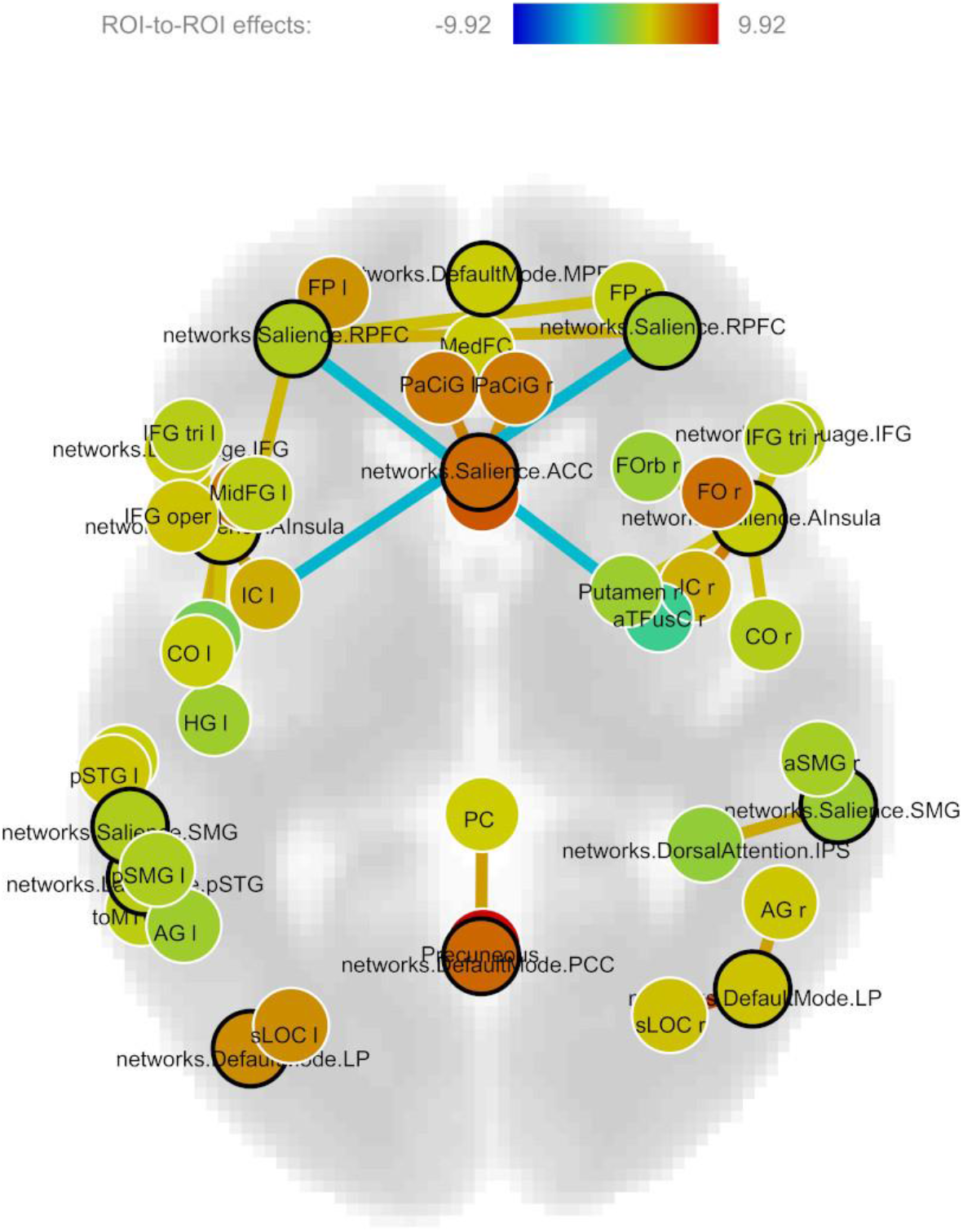
Functional connectivity for brain areas with statistically significant correlation with other regions of interest (ROI).

**Figure 3.**
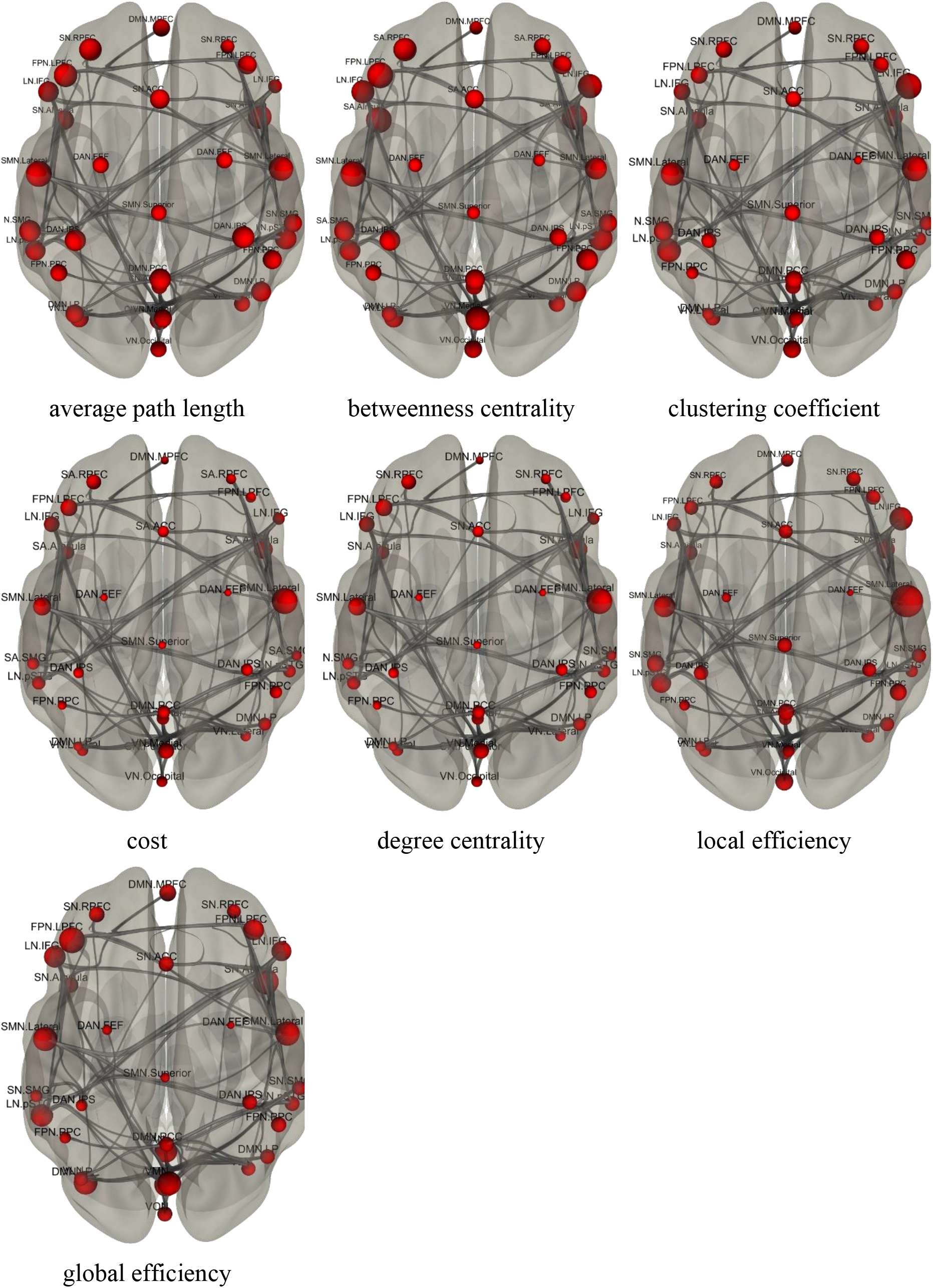
All of the graph parameters on one view.

**Figure 4.**
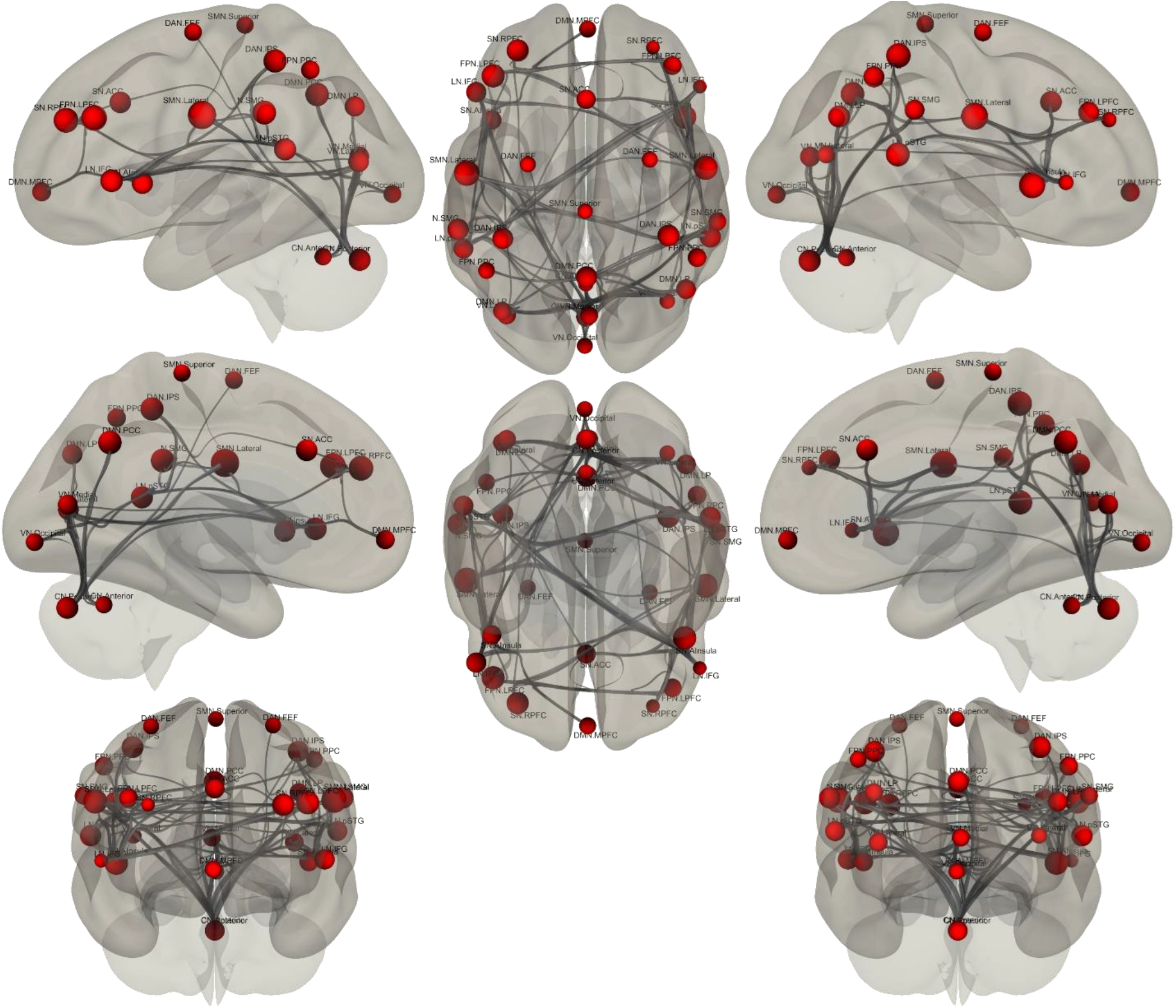
Graph parameter – *average path length* (the average distance from each node to any other node)

**Figure 5.**
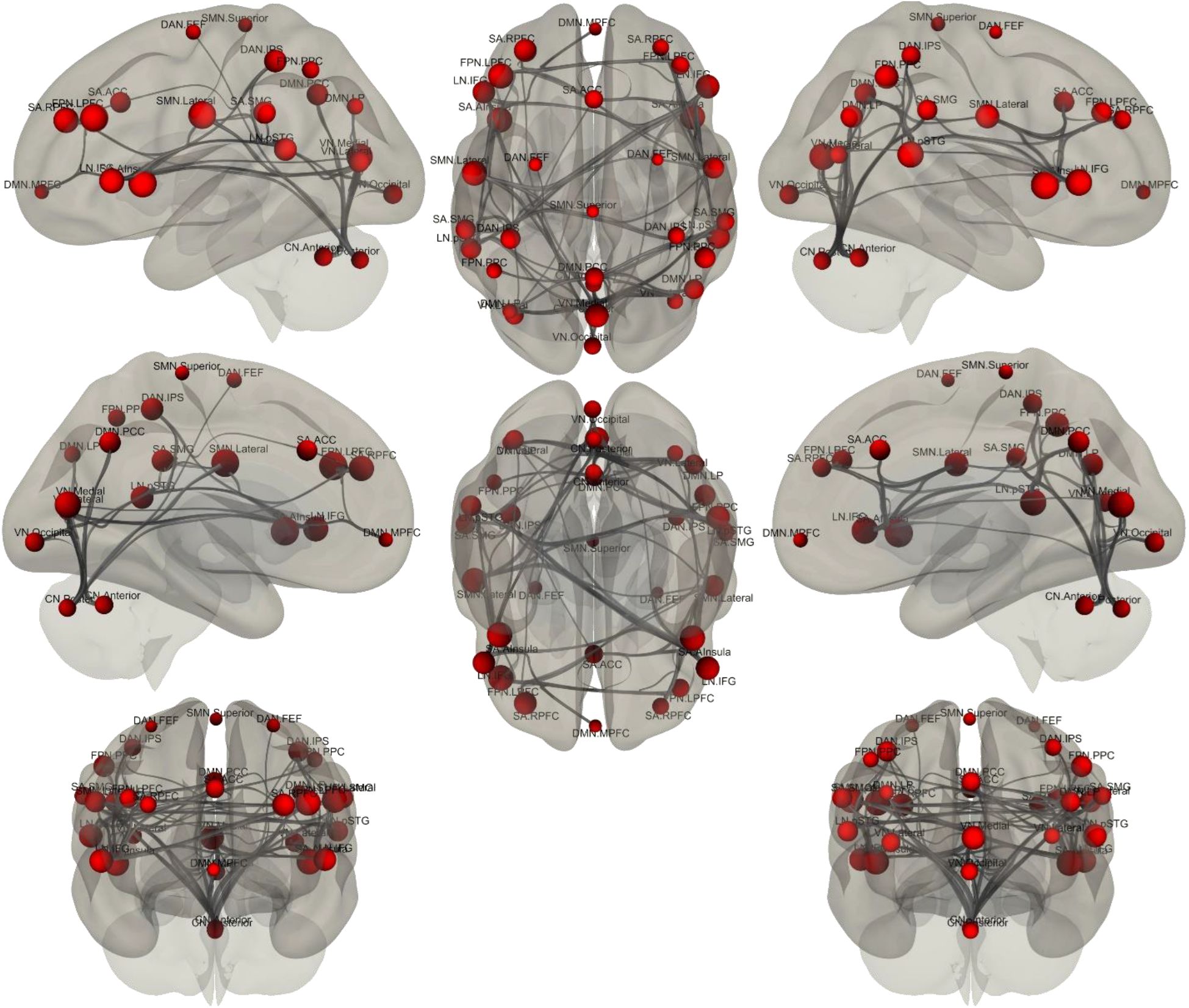
Graph parameter – *betweenness centrality* (the proportion of shortest paths between all node pairs in the network that pass through a given index node)

**Figure 6.**
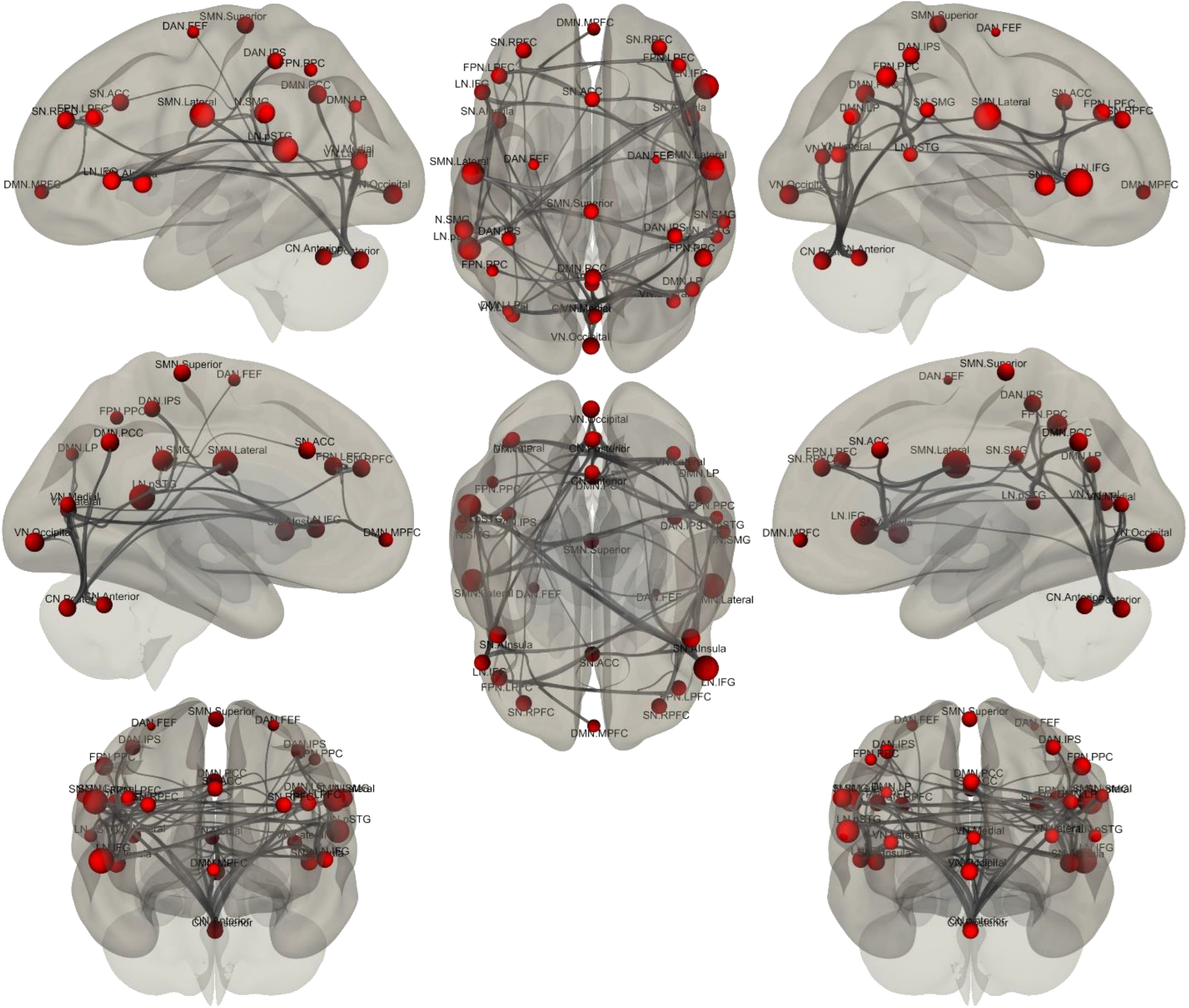
Graph parameter – *clustering coefficient* (the proportion of ROIs that have connectivity with a particular ROI that also have connectivity with each other)

**Figure 7.**
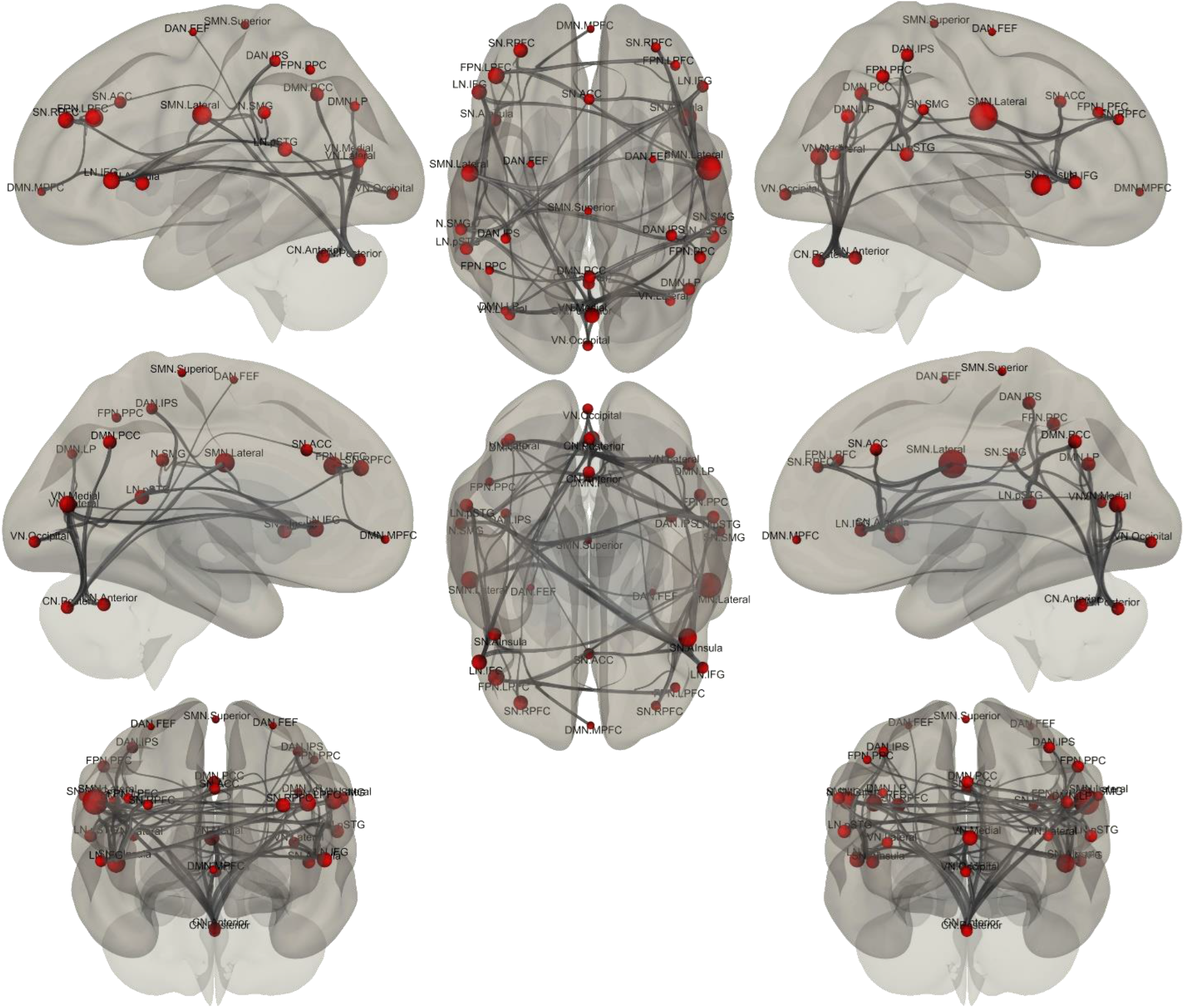
Graph parameter – *cost* (the ratio of the existing number of edges to the number of all possible edges in the network)

**Figure 8.**
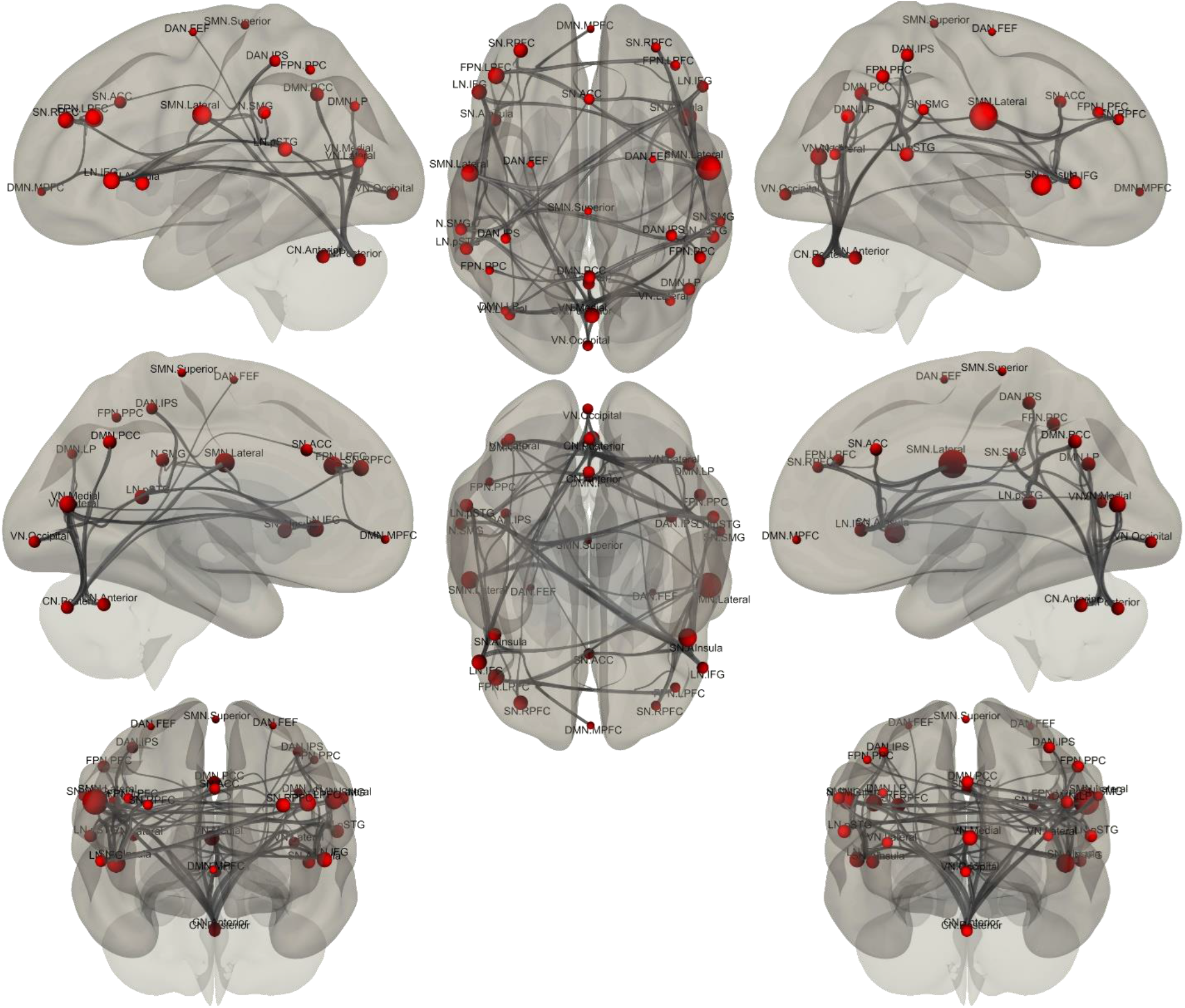
Graph parameter – *degree centrality* (the number of edges that connect a node to the rest of the network)

**Figure 9.**
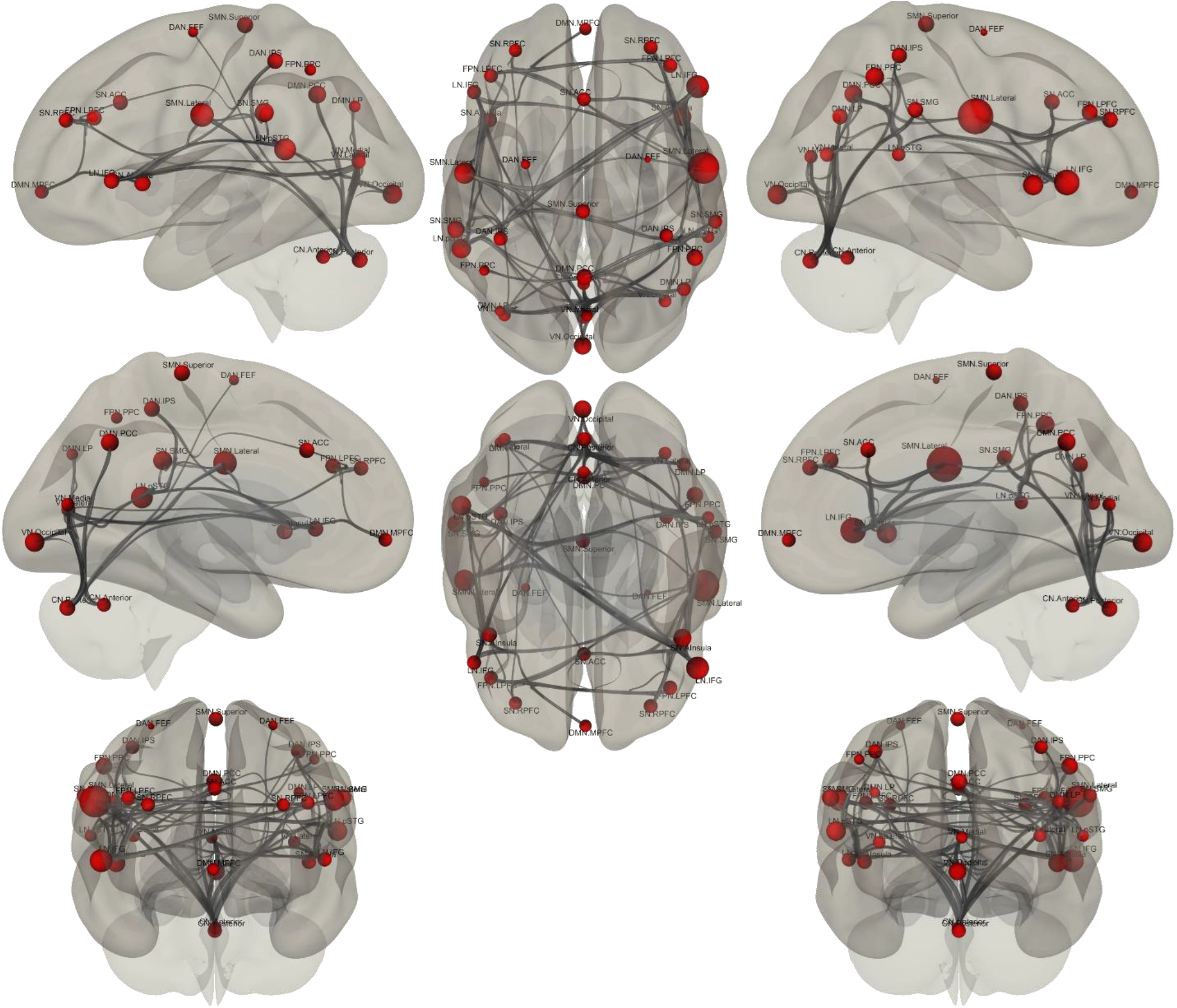
Graph parameter – *local efficiency* (the network ability in transmitting information at the local level)

**Figure 10.**
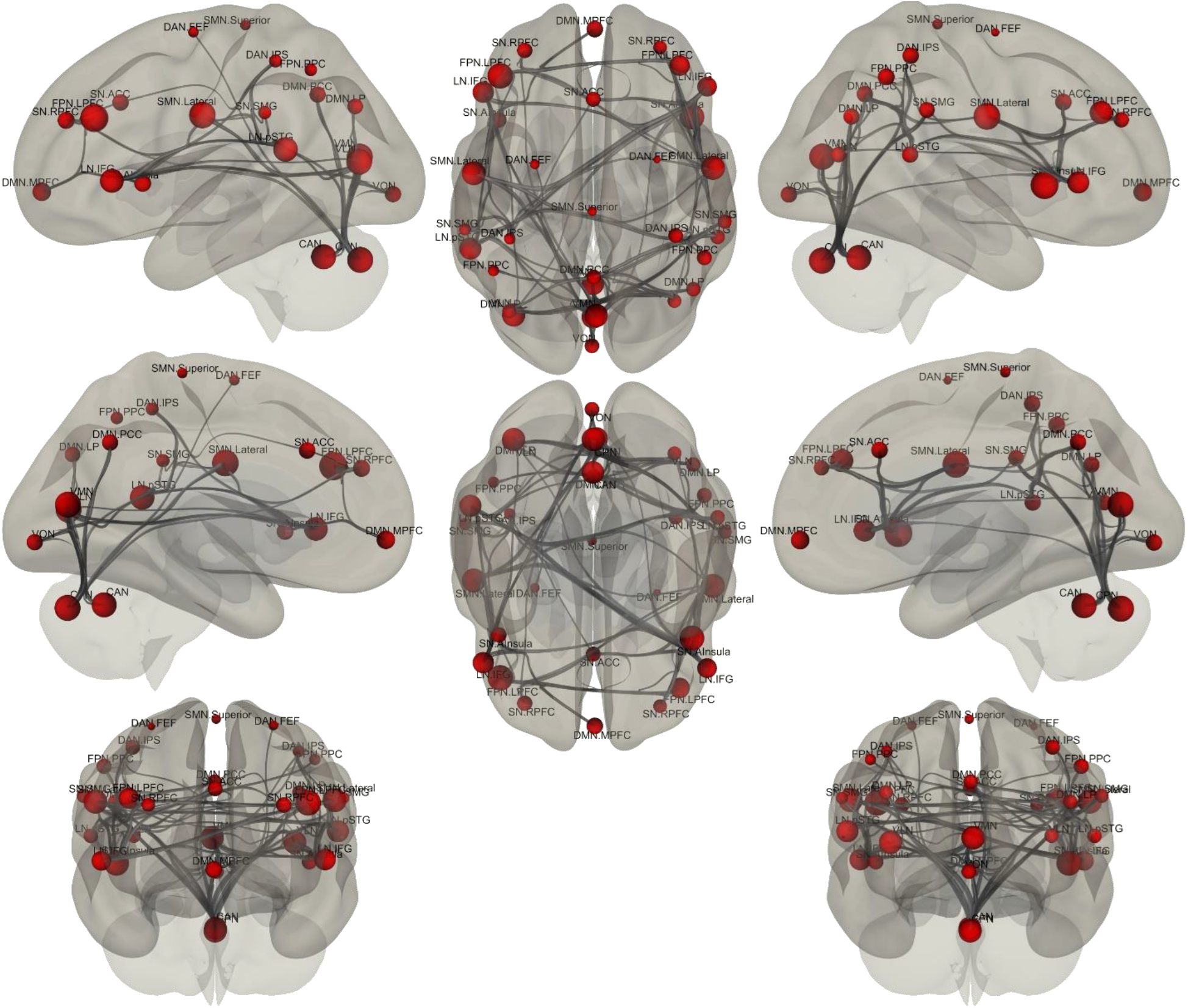
Graph parameter – *global efficiency* (the average inverse shortest path length in the network; this parameter is inversely related to the path length)

**Figure 11.**
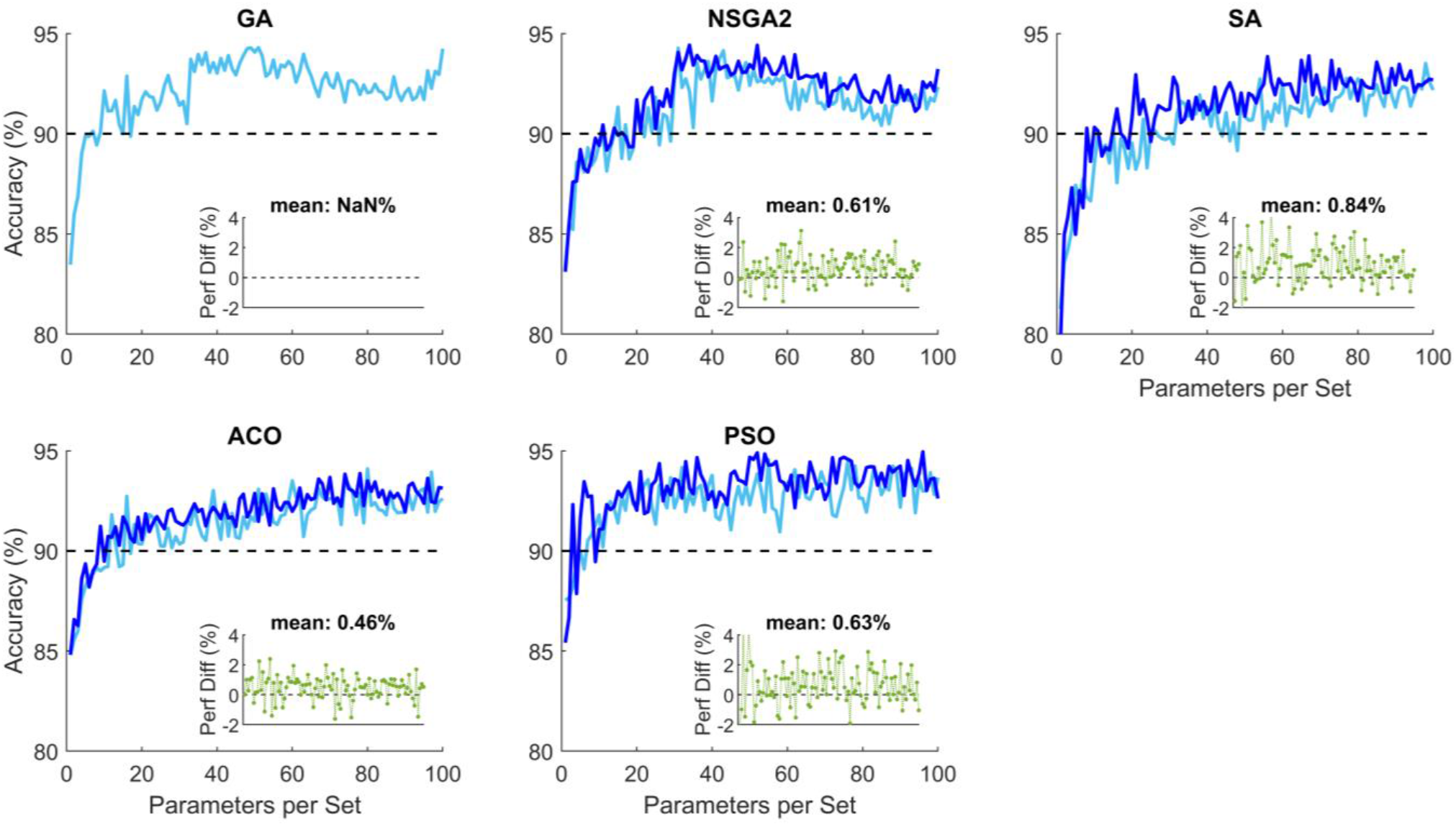
Comparison of classification performance for 200 repetitions (light blue) and 500 repetitions (dark blue) for different optimization algorithms per parameter set. The subplots show the difference between 200 and 500 repetitions, showing small superior performance for 500 repetitions. This is an indication that the algorithms converted within the first 200 repetitions.

**Supplementary Table 1.**
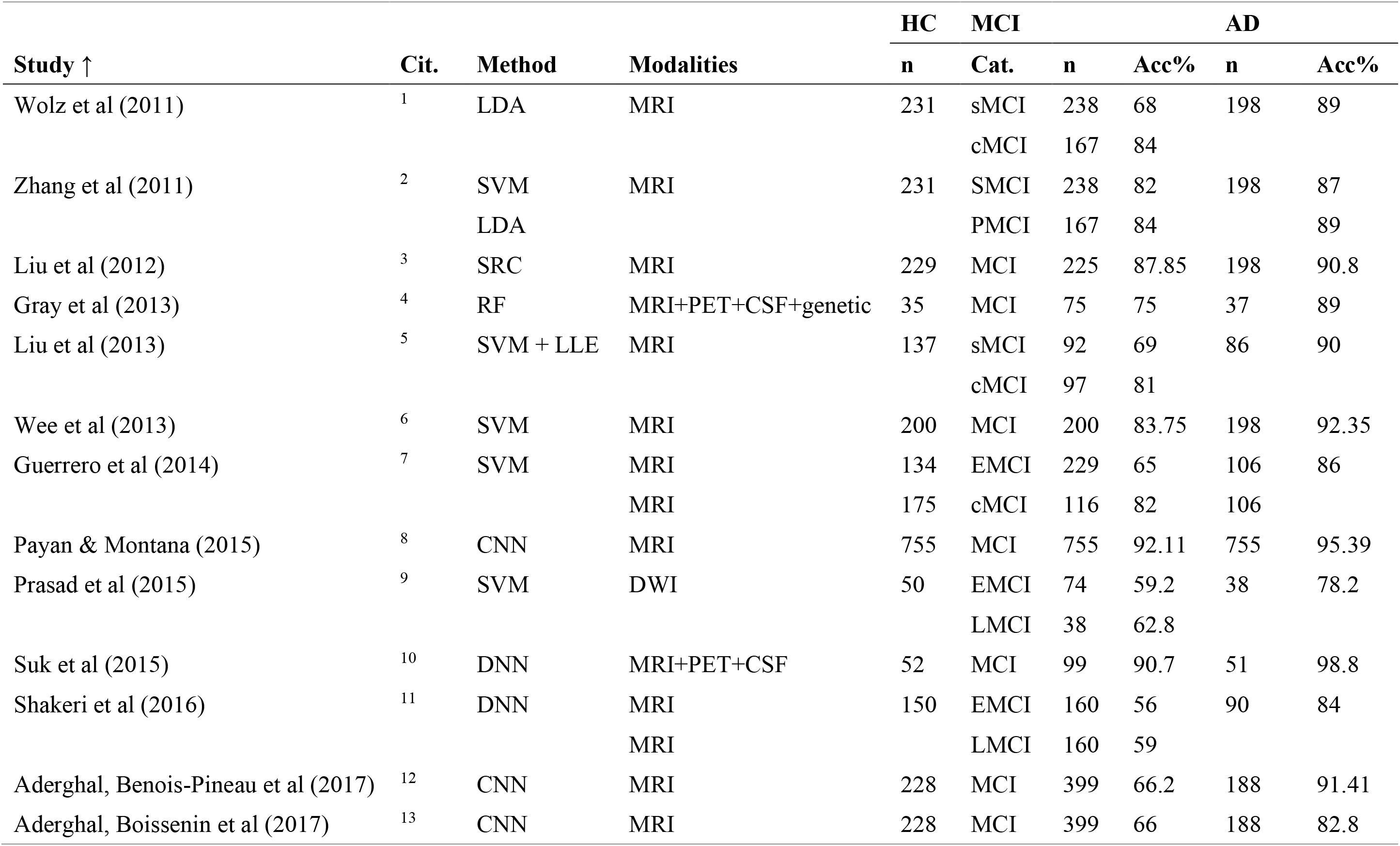
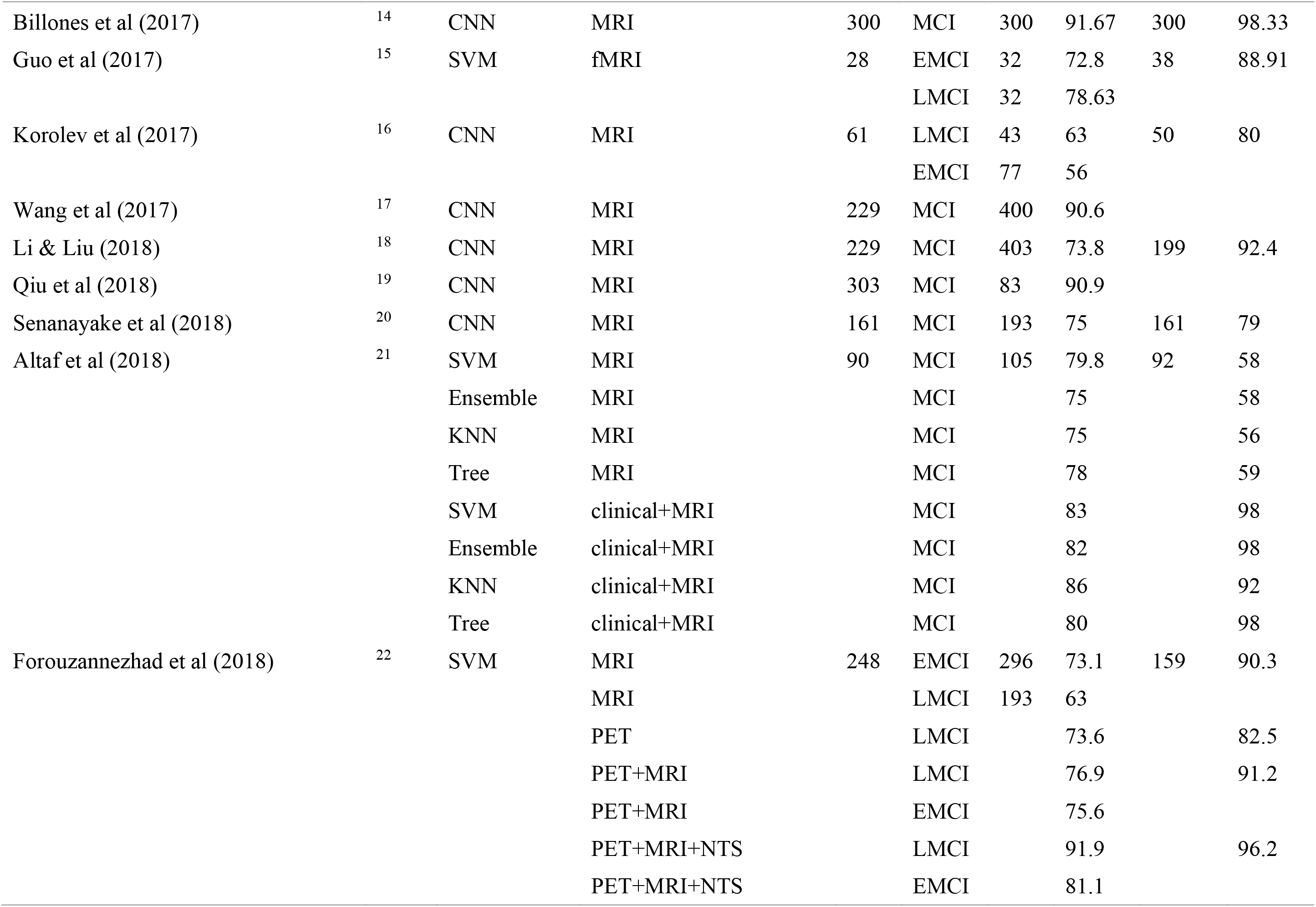
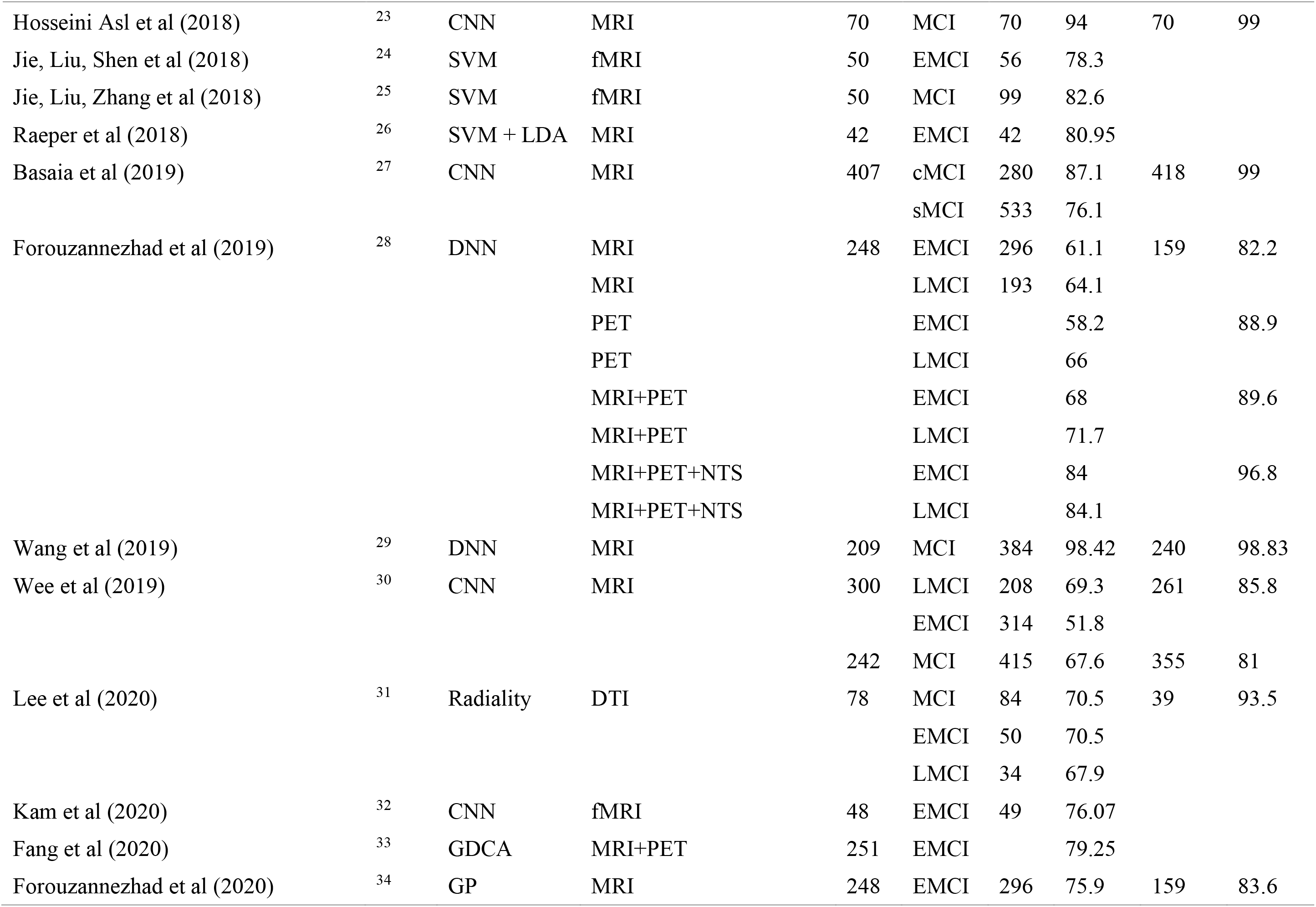
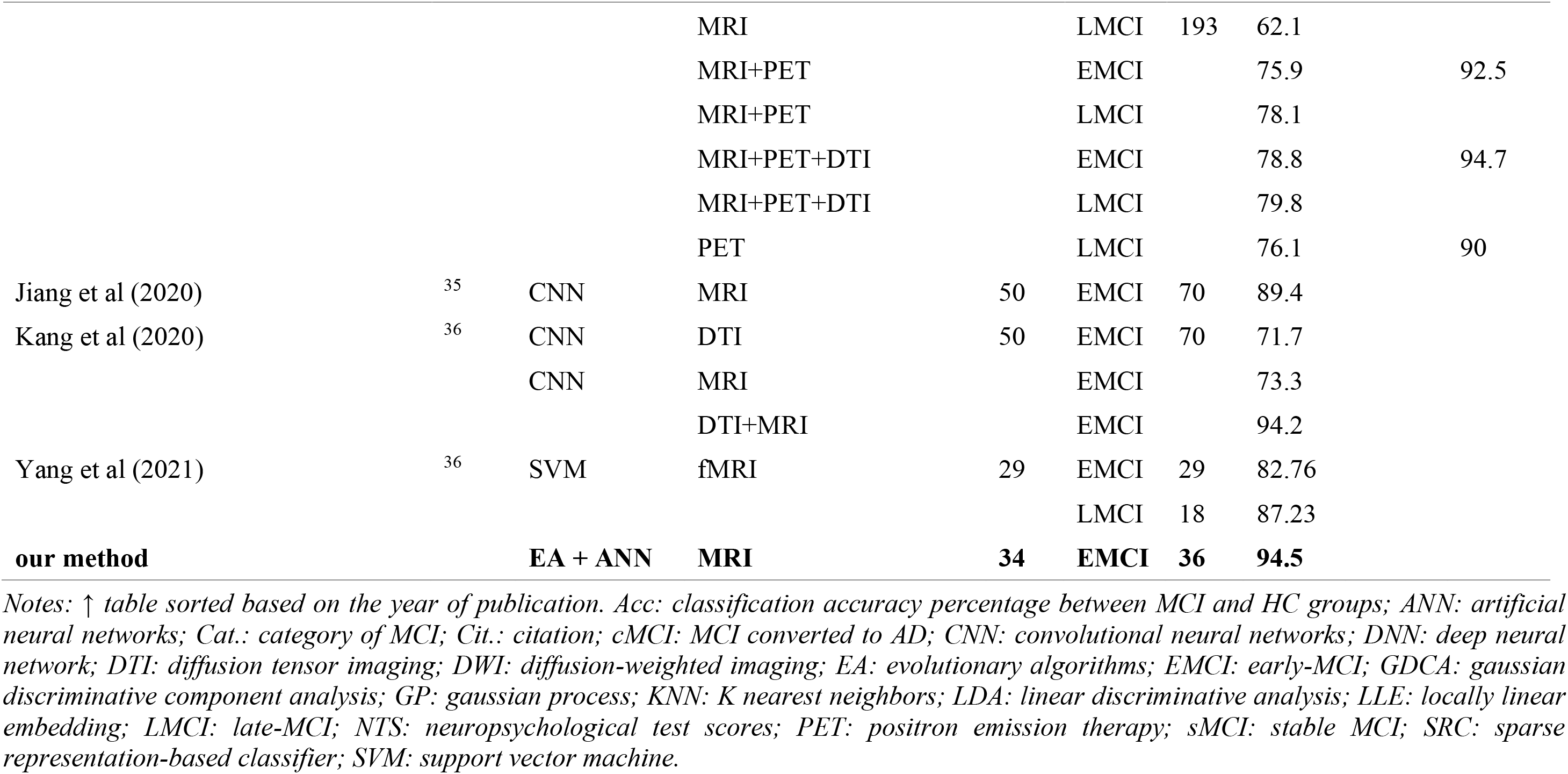
Summary of the studies aiming at categorization of healthy (HC), mild cognitive impairment (MCI) and Alzheimer’s disease (AD) using different biomarkers and classification methods based on Table 1.

## References

1. Scheltens, P. et al. Alzheimer’s disease. The Lancet 388, 505–517 (2016).

2. Nichols, E. et al. Global, regional, and national burden of Alzheimer’s disease and other dementias, 1990–2016: a systematic analysis for the Global Burden of Disease Study 2016. The Lancet Neurology 18, 88–106 (2019).

3. Braak, H. & Braak, E. Neuropathological stageing of Alzheimer-related changes. Acta Neuropathologica 82, 239–259 (1991).

4. Edmonds, E. C. et al. Early versus late MCI: Improved MCI staging using a neuropsychological approach. Alzheimer’s and Dementia 15, 699–708 (2019).

5. Platero, C. & Tobar, M. C. Predicting Alzheimer’s conversion in mild cognitive impairment patients using longitudinal neuroimaging and clinical markers. Brain Imaging and Behavior (2020) doi:10.1007/s11682-020-00366-8.

6. Petersen, R. C. et al. Practice guideline update summary: Mild cognitive impairment report of theguideline development, dissemination, and implementation. Neurology 90, 126–135 (2018).

7. Nestor, P. J., Scheltens, P. & Hodges, J. R. Advances in the early detection of Alzheimer’s disease. Nature Medicine 10, S34–S41 (2004).

8. Khazaee, A., Ebrahimzadeh, A. & Babajani-Feremi, A. Application of advanced machine learning methods on resting-state fMRI network for identification of mild cognitive impairment and Alzheimer’s disease. Brain Imaging and Behavior 10, 799–817 (2016).

9. Zhou, Q. et al. An Optimal Decisional Space for the Classification of Alzheimer’s Disease and Mild Cognitive Impairment. IEEE Transactions on Biomedical Engineering 61, 2245–2253 (2014).

10. Petersen, R. C. et al. Mild cognitive impairment: A concept in evolution. Journal of Internal Medicine 275, 214–228 (2014).

11. de Marco, M., Beltrachini, L., Biancardi, A., Frangi, A. F. & Venneri, A. Machine-learning Support to Individual Diagnosis of Mild Cognitive Impairment Using Multimodal MRI and Cognitive Assessments. Alzheimer Disease & Associated Disorders 31, 278–286 (2017).

12. Wang, B. et al. Early Stage Identification of Alzheimer’s Disease Using a Two-stage Ensemble Classifier. Current Bioinformatics 13, 529–535 (2018).

13. Zhang, T. et al. Classification of early and late mild cognitive impairment using functional brain network of resting-state fMRI. Frontiers in Psychiatry 10, 1–16 (2019).

14. Hojjati, S. H., Ebrahimzadeh, A. & Babajani-Feremi, A. Identification of the early stage of alzheimer’s disease using structural mri and resting-state fmri. Frontiers in Neurology 10, 1–12 (2019).

15. Bature, F., Guinn, B. A., Pang, D. & Pappas, Y. Signs and symptoms preceding the diagnosis of Alzheimer’s disease: A systematic scoping review of literature from 1937 to 2016. BMJ Open 7, (2017).

16. Reitz, C. & Mayeux, R. Alzheimer disease: Epidemiology, diagnostic criteria, risk factors and biomarkers. Biochemical Pharmacology 88, 640–651 (2014).

17. Dubois, B. et al. Preclinical Alzheimer’s disease: Definition, natural history, and diagnostic criteria. Alzheimer’s and Dementia vol. 12 (2016).

18. Tan, C. C., Yu, J. T. & Tan, L. Biomarkers for preclinical alzheimer’s disease. Journal of Alzheimer’s Disease 42, 1051–1069 (2014).

19. Schmand, B., Huizenga, H. M. & van Gool, W. A. Meta-analysis of CSF and MRI biomarkers for detecting preclinical Alzheimer’s disease. Psychological Medicine 40, 135–145 (2010).

20. Frisoni, G. B. et al. Strategic roadmap for an early diagnosis of Alzheimer’s disease based on biomarkers. The Lancet Neurology 16, 661–676 (2017).

21. Mueller, S. G. et al. Hippocampal atrophy patterns in mild cognitive impairment and alzheimer’s disease. Human Brain Mapping 31, 1339–1347 (2010).

22. Zamani, J., Sadr, A. & Javadi, A. A Large-scale Comparison of Cortical and Subcortical Structural Segmentation Methods in Alzheimer’ s Disease: a Statistical Approach. bioRxiv (2020) doi:10.1101/2020.08.18.256321.

23. Pini, L. et al. Brain atrophy in Alzheimer’s Disease and aging. Ageing Research Reviews 30, 25–48 (2016).

24. Frisoni, G. B., Fox, N. C., Jack, C. R., Scheltens, P. & Thompson, P. M. The clinical use of structural MRI in Alzheimer disease. Nature Reviews Neurology 6, 67–77 (2010).

25. Yassa, M. A. et al. High-resolution structural and functional MRI of hippocampal CA3 and dentate gyrus in patients with amnestic Mild Cognitive Impairment. NeuroImage 51, 1242–1252 (2010).

26. Sperling, R. The potential of functional MRI as a biomarker in early Alzheimer’s disease. Neurobiology of Aging 32, S37–S43 (2011).

27. Wierenga, C. E. & Bondi, M. W. Use of functional magnetic resonance imaging in the early identification of Alzheimer’s disease. Neuropsychology Review 17, 127–143 (2007).

28. Pievani, M., de Haan, W., Wu, T., Seeley, W. W. & Frisoni, G. B. Functional network disruption in the degenerative dementias. The Lancet Neurology 10, 829–843 (2011).

29. Teipel, S. et al. Multimodal imaging in Alzheimer’s disease: Validity and usefulness for early detection. The Lancet Neurology 14, 1037–1053 (2015).

30. Lee, M. H., Smyser, C. D. & Shimony, J. S. Resting-state fMRI: A review of methods and clinical applications. American Journal of Neuroradiology 34, 1866–1872 (2013).

31. Vemuri, P., Jones, D. T. & Jack, C. R. Resting state functional MRI in Alzheimer’s disease. Alzheimer’s Research and Therapy 4, 1–9 (2012).

32. Fox, M. D. & Greicius, M. Clinical applications of resting state functional connectivity. Frontiers in Systems Neuroscience 4, (2010).

33. Greicius, M. D., Krasnow, B., Reiss, A. L. & Menon, V. Functional connectivity in the resting brain: A network analysis of the default mode hypothesis. Proceedings of the National Academy of Sciences of the United States of America 100, 253–258 (2003).

34. Sheline, Y. I. & Raichle, M. E. Resting state functional connectivity in preclinical Alzheimer’s disease. Biological Psychiatry 74, 340–347 (2013).

35. Zhan, Y. et al. Longitudinal Study of Impaired Intra- and Inter-Network Brain Connectivity in Subjects at High Risk for Alzheimer’s Disease. Journal of Alzheimer’s Disease 52, 913–927 (2016).

36. Zhang, H. Y. et al. Resting brain connectivity: Changes during the progress of Alzheimer disease. Radiology 256, 598–606 (2010).

37. Zhou, J. et al. Divergent network connectivity changes in behavioural variant frontotemporal dementia and Alzheimer’s disease. Brain 133, 1352–1367 (2010).

38. Dennis, E. L. & Thompson, P. M. Functional brain connectivity using fMRI in aging and Alzheimer’s disease. Neuropsychology Review 24, 49–62 (2014).

39. Bullmore, E. & Sporns, O. Complex brain networks: Graph theoretical analysis of structural and functional systems. Nature Reviews Neuroscience 10, 186–198 (2009).

40. Rubinov, M. & Sporns, O. Complex network measures of brain connectivity: Uses and interpretations. NeuroImage 52, 1059–1069 (2010).

41. van den Heuvel, M. P. & Sporns, O. Network hubs in the human brain. Trends in Cognitive Sciences 17, 683–696 (2013).

42. Farahani, F. v., Karwowski, W. & Lighthall, N. R. Application of graph theory for identifying connectivity patterns in human brain networks: A systematic review. Frontiers in Neuroscience 13, 1–27 (2019).

43. He, Y. & Evans, A. Graph theoretical modeling of brain connectivity. Current Opinion in Neurology 23, 341–350 (2010).

44. Bassett, D. S. & Bullmore, E. T. Human brain networks in health and disease. Current Opinion in Neurology 22, 340–347 (2009).

45. Stam, C. J. Modern network science of neurological disorders. Nature Reviews Neuroscience 15, 683–695 (2014).

46. Tijms, B. M. et al. Alzheimer’s disease: connecting findings from graph theoretical studies of brain networks. Neurobiology of Aging 34, 2023–2036 (2013).

47. Brier, M. R. et al. Functional connectivity and graph theory in preclinical Alzheimer’s disease. Neurobiology of Aging 35, 757–768 (2014).

48. Dai, Z. et al. Identifying and mapping connectivity patterns of brain network hubs in Alzheimer’s disease. Cerebral Cortex 25, 3723–3742 (2015).

49. Khazaee, A., Ebrahimzadeh, A. & Babajani-Feremi, A. Identifying patients with Alzheimer’s disease using resting-state fMRI and graph theory. Clinical Neurophysiology 126, 2132–2141 (2015).

50. Hojjati, S. H., Ebrahimzadeh, A., Khazaee, A. & Babajani-Feremi, A. Predicting conversion from MCI to AD using resting-state fMRI, graph theoretical approach and SVM. Journal of Neuroscience Methods 282, 69–80 (2017).

51. Khazaee, A., Ebrahimzadeh, A. & Babajani-Feremi, A. Classification of patients with MCI and AD from healthy controls using directed graph measures of resting-state fMRI. Behavioural Brain Research 322, 339–350 (2017).

52. Blum, A. L. & Langley, P. Selection of relevant features and examples in machine learning. Artificial Intelligence 97, 245–271 (1997).

53. Reunanen, J. Overfitting in making comparisons between variable selection methods. Journal of Machine Learning Research 3, 1371–1382 (2003).

54. John, G. H., Kohavi, R. & Pfleger, K. Irrelevant Features and the Subset Selection Problem. in Machine Learning Proceedings 1994 121–129 (Elsevier, 1994). doi:10.1016/B978-1-55860-335-6.50023-4.

55. Chu, C., Hsu, A. L., Chou, K. H., Bandettini, P. & Lin, C. P. Does feature selection improve classification accuracy? Impact of sample size and feature selection on classification using anatomical magnetic resonance images. NeuroImage 60, 59–70 (2012).

56. Xue, B., Zhang, M., Browne, W. N. & Yao, X. A Survey on Evolutionary Computation Approaches to Feature Selection. IEEE Transactions on Evolutionary Computation 20, 606–626 (2016).

57. Jaderberg, M. et al. Population based training of neural networks. arXiv (2017).

58. del Ser, J. et al. Bio-inspired computation: Where we stand and what’s next. Swarm and Evolutionary Computation 48, 220–250 (2019).

59. Zawbaa, H. M., Emary, E., Grosan, C. & Snasel, V. Large-dimensionality small-instance set feature selection: A hybrid bio-inspired heuristic approach. Swarm and Evolutionary Computation 42, 29–42 (2018).

60. Gu, S., Cheng, R. & Jin, Y. Feature selection for high-dimensional classification using a competitive swarm optimizer. Soft Computing 22, 811–822 (2018).

61. Sharma, M., Pradhyumna, S. P., Goyal, S. & Singh, K. Machine Learning and Evolutionary Algorithms for the Diagnosis and Detection of Alzheimer’s Disease. in Data Analytics and Management. Lecture Notes on Data Engineering and Communications Technologies (eds. Khanna, A., Gupta, D., Z, P., S, B. & O, C.) 229–250 (Springer, 2021). doi:10.1007/978-981-15-8335-3_20.

62. Dessouky, M. & Elrashidy, M. Feature Extraction of the Alzheimer’s Disease Images Using Different Optimization Algorithms. Journal of Alzheimer’s Disease & Parkinsonism 6, (2016).

63. Johnson, P. et al. Genetic algorithm with logistic regression for prediction of progression to Alzheimer’s disease. BMC Bioinformatics 15, 1–14 (2014).

64. Kroll, J. P., Eickhoff, S. B., Hoffstaedter, F. & Patil, K. R. Evolving complex yet interpretable representations: Application to Alzheimer’s diagnosis and prognosis. 2020 IEEE Congress on Evolutionary Computation, CEC 2020 - Conference Proceedings (2020) doi:10.1109/CEC48606.2020.9185843.

65. Bi, X. A., Xu, Q., Luo, X., Sun, Q. & Wang, Z. Analysis of progression toward Alzheimer’s disease based on evolutionary weighted random support vector machine cluster. Frontiers in Neuroscience 12, 1–11 (2018).

66. Jack, C. R. et al. Introduction to the recommendations from the National Institute on Aging-Alzheimer’s Association workgroups on diagnostic guidelines for Alzheimer’s disease. Alzheimer’s and Dementia 7, 257–262 (2011).

67. Jack, C. R. et al. Update on the Magnetic Resonance Imaging core of the Alzheimer’s Disease Neuroimaging Initiative. Alzheimer’s and Dementia 6, 212–220 (2010).

68. Jack, C. R. et al. The Alzheimer’s Disease Neuroimaging Initiative (ADNI): MRI methods. Journal of Magnetic Resonance Imaging 27, 685–691 (2008).

69. Aisen, P. S., Petersen, R. C., Donohue, M. & Weiner, M. W. Alzheimer’s Disease Neuroimaging Initiative 2 Clinical Core: Progress and plans. Alzheimer’s and Dementia 11, 734–739 (2015).

70. Nieto-Castanon, A. Handbook of fcMRI methods in CONN. (2020).

71. Whitfield-Gabrieli, S. & Nieto-Castanon, A. Conn: A Functional Connectivity Toolbox for Correlated and Anticorrelated Brain Networks. Brain Connectivity 2, 125–141 (2012).

72. Biswal, B., Zerrin Yetkin, F., Haughton, V. M. & Hyde, J. S. Functional connectivity in the motor cortex of resting human brain using echo-planar MRI. Magnetic Resonance in Medicine 34, 537–541 (1995).

73. Damoiseaux, J. S. et al. Reduced resting-state brain activity in the “default network” in normal aging. Cerebral Cortex 18, 1856–1864 (2008).

74. Fox, M. D. et al. The human brain is intrinsically organized into dynamic, anticorrelated functional networks. Proceedings of the National Academy of Sciences of the United States of America 102, 9673–9678 (2005).

75. Tzourio-Mazoyer, N. et al. Automated Anatomical Labeling of Activations in SPM Using a Macroscopic Anatomical Parcellation of the MNI MRI Single-Subject Brain. NeuroImage 15, 273–289 (2002).

76. Jafri, M. J., Pearlson, G. D., Stevens, M. & Calhoun, V. D. A method for functional network connectivity among spatially independent resting-state components in schizophrenia. NeuroImage 39, 1666–1681 (2008).

77. Joel, S. E., Caffo, B. S., van Zijl, P. C. M. & Pekar, J. J. On the relationship between seed-based and ICA-based measures of functional connectivity. Magnetic Resonance in Medicine 66, 644–657 (2011).

78. delEtoile, J. & Adeli, H. Graph Theory and Brain Connectivity in Alzheimer’s Disease. Neuroscientist 23, 616–626 (2017).

79. Fornito, A., Andrew, Z. & Edward, B. Fundamentals of brain network analysis. (Academic Press, 2016).

80. Tsai, C. F., Eberle, W. & Chu, C. Y. Genetic algorithms in feature and instance selection. Knowledge-Based Systems 39, 240–247 (2013).

81. Goldenberg, D. E. Genetic algorithms in search, optimization and machine learning. (Addison Wesley, 1989).

82. Vandewater, L., Brusic, V., Wilson, W., Macaulay, L. & Zhang, P. An adaptive genetic algorithm for selection of blood-based biomarkers for prediction of Alzheimer’s disease progression. BMC Bioinformatics 16, 1–10 (2015).

83. Deb, K., Pratap, A., Agarwal, S. & Meyarivan, T. A fast and elitist multiobjective genetic algorithm: NSGA-II. IEEE Transactions on Evolutionary Computation 6, 182–197 (2002).

84. Dorigo, M., Caro, G. di & Gambardella, L. M. Ant Algorithms for Discrete Optimization. Artificial Life 5, 137–172 (1999).

85. Akhtar, A. Evolution of Ant Colony Optimization Algorithm — A Brief Literature Review. arXiv (2019).

86. Kalami Heris, S. M. & Khaloozadeh, H. Ant Colony Estimator: An intelligent particle filter based on ACO ℝ. Engineering Applications of Artificial Intelligence 28, 78–85 (2014).

87. Kirkpatrick, S., Gelatt, C. D. & Vecchi, M. P. Optimization by Simulated Annealing. Science 220, 671–680 (1983).

88. Anily, S. & Federgruen, A. Simulated Annealing Methods With General Acceptance Probabilities. Journal of Applied Probability 24, 657–667 (1987).

89. Bertsimas, D. & Tsitsiklis, J. Simulated annealing. Statistical Science 8, 10–15 (1993).

90. Kennedy, J. & Eberhart, R. Particle swarm optimization. in Proceedings of ICNN’95 - International Conference on Neural Networks vol. 4 1942–1948 (IEEE, 1995).

91. Wang, X., Yang, J., Teng, X., Xia, W. & Jensen, R. Feature selection based on rough sets and particle swarm optimization. Pattern Recognition Letters 28, 459–471 (2007).

92. Team, Y. Particle swarm optimization in MATLAB. (2015).

93. Lv, C. et al. Levenberg-marquardt backpropagation training of multilayer neural networks for state estimation of a safety-critical cyber-physical system. IEEE Transactions on Industrial Informatics 14, 3436–3446 (2018).

94. de Rubio, J. J. Stability Analysis of the Modified Levenberg-Marquardt Algorithm for the Artificial Neural Network Training. IEEE Transactions on Neural Networks and Learning Systems 1–15 (2020) doi:10.1109/TNNLS.2020.3015200.

95. Hagan, M. T. & Menhaj, M. B. Training feedforward networks with the Marquardt algorithm. IEEE Transactions on Neural Networks 5, 989–993 (1994).

96. Wen, J. et al. Convolutional neural networks for classification of Alzheimer’s disease: Overview and reproducible evaluation. Medical Image Analysis 63, 101694 (2020).

97. Payan, A. & Montana, G. Predicting Alzheimer’s disease a neuroimaging study with 3D convolutional neural networks. ICPRAM 2015 - 4th International Conference on Pattern Recognition Applications and Methods, Proceedings 2, 355–362 (2015).

98. Wang, H. et al. Ensemble of 3D densely connected convolutional network for diagnosis of mild cognitive impairment and Alzheimer’s disease. Neurocomputing 333, 145–156 (2019).

99. Billones, C. D., Demetria, O. J. L. D., Hostallero, D. E. D. & Naval, P. C. DemNet: A Convolutional Neural Network for the detection of Alzheimer’s Disease and Mild Cognitive Impairment. IEEE Region 10 Annual International Conference, Proceedings/TENCON 3724–3727 (2017) doi:10.1109/TENCON.2016.7848755.

100. Wang, S., Shen, Y., Chen, W., Xiao, T. & Hu, J. Automatic Recognition of Mild Cognitive Impairment from MRI Images Using Expedited Convolutional Neural Networks. in Lecture Notes in Computer Science (including subseries Lecture Notes in Artificial Intelligence and Lecture Notes in Bioinformatics) vol. 10613 LNCS 373–380 (2017).

101. Qiu, S. et al. Fusion of deep learning models of MRI scans, Mini–Mental State Examination, and logical memory test enhances diagnosis of mild cognitive impairment. Alzheimer’s and Dementia: Diagnosis, Assessment and Disease Monitoring 10, 737–749 (2018).

102. Forouzannezhad, P., Abbaspour, A., Cabrerizo, M. & Adjouadi, M. Early Diagnosis of Mild Cognitive Impairment Using Random Forest Feature Selection. in 2018 IEEE Biomedical Circuits and Systems Conference (BioCAS) vol. 53 1–4 (IEEE, 2018).

103. Wolz, R. et al. Multi-method analysis of MRI images in early diagnostics of Alzheimer’s disease. PLoS ONE 6, 1–9 (2011).

104. Zhang, D., Wang, Y., Zhou, L., Yuan, H. & Shen, D. Multimodal classification of Alzheimer’s disease and mild cognitive impairment. NeuroImage 55, 856–867 (2011).

105. Liu, M., Zhang, D. & Shen, D. Ensemble sparse classification of Alzheimer’s disease. NeuroImage 60, 1106–1116 (2012).

106. Gray, K. R., Aljabar, P., Heckemann, R. A., Hammers, A. & Rueckert, D. Random forest-based similarity measures for multi-modal classification of Alzheimer’s disease. NeuroImage 65, 167–175 (2013).

107. Liu, X., Tosun, D., Weiner, M. W. & Schuff, N. Locally linear embedding (LLE) for MRI based Alzheimer’s disease classification. NeuroImage 83, 148–157 (2013).

108. Wee, C. Y., Yap, P. T. & Shen, D. Prediction of Alzheimer’s disease and mild cognitive impairment using cortical morphological patterns. Human Brain Mapping 34, 3411–3425 (2013).

109. Guerrero, R., Wolz, R., Rao, A. W. & Rueckert, D. Manifold population modeling as a neuro-imaging biomarker: Application to ADNI and ADNI-GO. NeuroImage 94, 275–286 (2014).

110. Prasad, G., Joshi, S. H., Nir, T. M., Toga, A. W. & Thompson, P. M. Brain connectivity and novel network measures for Alzheimer’s disease classification. Neurobiology of Aging 36, S121–S131 (2015).

111. Suk, H. il, Lee, S. W. & Shen, D. Latent feature representation with stacked auto-encoder for AD/MCI diagnosis. Brain Structure and Function 220, 841–859 (2015).

112. Shakeri, M., Lombaert, H., Tripathi, S. & Kadoury, S. Deep spectral-based shape features for Alzheimer’s disease classification. Lecture Notes in Computer Science (including subseries Lecture Notes in Artificial Intelligence and Lecture Notes in Bioinformatics) 10126 LNCS, 15–24 (2016).

113. Aderghal, K., Benois-Pineau, J., Afdel, K. & Gwenaëlle, C. FuseMe: Classification of sMRI images by fusion of deep CNNs in 2D+e projections. ACM International Conference Proceeding Series Part F1301, (2017).

114. Aderghal, K., Boissenin, M., Benois-Pineau, J., Catheline, G. & Afdel, K. Classification of sMRI for AD Diagnosis with Convolutional Neuronal Networks: A Pilot 2-D+ ϵ Study on ADNI. in MultiMedia Modeling, Lecture Notes in Computer Science (eds. Amsaleg, L., Gu ðmundsson, G., Gurrin, C., Jónsson, B. & Satoh, S.) 690–701 (Springer International Publishing, 2017). doi:10.1007/978-3-319-51811-4_56.

115. Guo, H., Zhang, F., Chen, J., Xu, Y. & Xiang, J. Machine learning classification combining multiple features of a hyper-network of fMRI data in Alzheimer’s disease. Frontiers in Neuroscience 11, 1–22 (2017).

116. Korolev, S., Safiullin, A., Belyaev, M. & Dodonova, Y. Residual and plain convolutional neural networks for 3D brain MRI classification. arXiv 835–838 (2017).

117. Li, F. & Liu, M. Alzheimer’s disease diagnosis based on multiple cluster dense convolutional networks. Computerized Medical Imaging and Graphics 70, 101–110 (2018).

118. Senanayake, U., Sowmya, A. & Dawes, L. Deep fusion pipeline for mild cognitive impairment diagnosis. in 2018 IEEE 15th International Symposium on Biomedical Imaging (ISBI 2018) 1394–1997 (IEEE, 2018). doi:10.1109/ISBI.2018.8363832.

119. Altaf, T., Anwar, S. M., Gul, N., Majeed, M. N. & Majid, M. Multi-class Alzheimer’s disease classification using image and clinical features. Biomedical Signal Processing and Control 43, 64–74 (2018).

120. Hosseini-Asl, E., Gimel’farb, G. & El-Baz, A. Alzheimer’s disease diagnostics by a deeply supervised adaptable 3D convolutional network. Frontiers in Biosciences 23, 584–596 (2018).

121. Jie, B., Liu, M. & Shen, D. Integration of temporal and spatial properties of dynamic connectivity networks for automatic diagnosis of brain disease. Medical Image Analysis 47, 81–94 (2018).

122. Jie, B., Liu, M., Zhang, D. & Shen, D. Sub-Network Kernels for Measuring Similarity of Brain Connectivity Networks in Disease Diagnosis. IEEE Transactions on Image Processing 27, 2340–2353 (2018).

123. Raeper, R., Lisowska, A. & Rekik, I. Cooperative Correlational and Discriminative Ensemble Classifier Learning for Early Dementia Diagnosis Using Morphological Brain Multiplexes. IEEE Access 6, 43830–43839 (2018).

124. Basaia, S. et al. Automated classification of Alzheimer’s disease and mild cognitive impairment using a single MRI and deep neural networks. NeuroImage: Clinical 21, 101645 (2019).

125. Forouzannezhad, P., Abbaspour, A., Li, C., Cabrerizo, M. & Adjouadi, M. A Deep Neural Network Approach for Early Diagnosis of Mild Cognitive Impairment Using Multiple Features. Proceedings - 17th IEEE International Conference on Machine Learning and Applications, ICMLA 2018 1341–1346 (2019) doi:10.1109/ICMLA.2018.00218.

126. Wee, C. Y. et al. Cortical graph neural network for AD and MCI diagnosis and transfer learning across populations. NeuroImage: Clinical 23, 101929 (2019).

127. Lee, P., Kim, H. R. & Jeong, Y. Detection of gray matter microstructural changes in Alzheimer’s disease continuum using fiber orientation. BMC Neurology 20, 1–10 (2020).

128. Kam, T. E., Zhang, H., Jiao, Z. & Shen, Di. Deep Learning of Static and Dynamic Brain Functional Networks for Early MCI Detection. IEEE Transactions on Medical Imaging 39, 478–487 (2020).

129. Fang, C. et al. Gaussian discriminative component analysis for early detection of Alzheimer’s disease: A supervised dimensionality reduction algorithm. Journal of Neuroscience Methods 344, 108856 (2020).

130. Forouzannezhad, P. et al. A Gaussian-based model for early detection of mild cognitive impairment using multimodal neuroimaging. Journal of Neuroscience Methods 333, 108544 (2020).

131. Jiang, J., Kang, L., Huang, J. & Zhang, T. Deep learning based mild cognitive impairment diagnosis using structure MR images. Neuroscience Letters 730, 134971 (2020).

132. Kang, L., Jiang, J., Huang, J. & Zhang, T. Identifying Early Mild Cognitive Impairment by Multi-Modality MRI-Based Deep Learning. Frontiers in Aging Neuroscience 12, 1–10 (2020).

133. Yang, P. et al. Fused Sparse Network Learning for Longitudinal Analysis of Mild Cognitive Impairment. IEEE Transactions on Cybernetics 51, 233–246 (2021).

134. Reyes, M. et al. On the Interpretability of Artificial Intelligence in Radiology: Challenges and Opportunities. Radiology: Artificial Intelligence 2, e190043 (2020).

135. Amorim, J. P., Abreu, P. H., Reyes, M. & Santos, J. Interpretability vs. Complexity: The Friction in Deep Neural Networks. Proceedings of the International Joint Conference on Neural Networks 1–7 (2020) doi:10.1109/IJCNN48605.2020.9206800.

136. Pereira, S. et al. Enhancing interpretability of automatically extracted machine learning features: application to a RBM-Random Forest system on brain lesion segmentation. Medical Image Analysis 44, 228–244 (2018).

137. Fan, F., Xiong, J. & Wang, G. On Interpretability of Artificial Neural Networks. arXiv (2020).

138. Beheshti, I., Demirel, H. & Matsuda, H. Classification of Alzheimer’s disease and prediction of mild cognitive impairment-to-Alzheimer’s conversion from structural magnetic resource imaging using feature ranking and a genetic algorithm. Computers in Biology and Medicine 83, 109–119 (2017).

139. Šprogar, M., Šprogar, M. & Colnarič, M. Autonomous evolutionary algorithm in medical data analysis. Computer Methods and Programs in Biomedicine 80, (2005).

140. Trunk, G. v. A Problem of Dimensionality: A Simple Example. IEEE Transactions on Pattern Analysis and Machine Intelligence PAMI-1, 306–307 (1979).

141. Hughes, G. F. On the Mean Accuracy of Statistical Pattern Recognizers. IEEE Transactions on Information Theory 14, 55–63 (1968).

142. Zollanvari, A., James, A. P. & Sameni, R. A Theoretical Analysis of the Peaking Phenomenon in Classification. Journal of Classification 37, 421–434 (2020).

143. Jain, A. K., Duin, R. P. W. & Mao, J. Statistical pattern recognition: A review. IEEE Transactions on Pattern Analysis and Machine Intelligence 22, 4–37 (2000).

## References

1. Amini, F. & Hu, G. A two-layer feature selection method using Genetic Algorithm and Elastic Net. Expert Systems with Applications 166, 114072 (2021).

2. Deb, K., Pratap, A., Agarwal, S. & Meyarivan, T. A fast and elitist multiobjective genetic algorithm: NSGA-II. IEEE Transactions on Evolutionary Computation 6, 182–197 (2002).

3. Srinivas, N. & Deb, K. Muiltiobjective Optimization Using Nondominated Sorting in Genetic Algorithms. Evolutionary Computation 2, 221–248 (1994).

4. Heris, S. M. K. & Khaloozadeh, H. Open-and closed-loop multiobjective optimal strategies for HIV therapy using NSGA-II. IEEE Transactions on Biomedical Engineering 58, 1678–1685 (2011).

5. Dang, V. Q. & Lam, C. NSC-NSGA2: Optimal search for finding multiple thresholds for nearest shrunken centroid. in 2013 IEEE International Conference on Bioinformatics and Biomedicine 367–372 (IEEE, 2013). doi:10.1109/BIBM.2013.6732520.

6. Kanan, H. R., Faez, K. & Taheri, S. M. Feature Selection Using Ant Colony Optimization (ACO): A New Method and Comparative Study in the Application of Face Recognition System. in Advances in Data Mining. Theoretical Aspects and Applications vol. 4597 LNCS 63–76 (Springer Berlin Heidelberg, 2007).

7. Lin, S. W., Lee, Z. J., Chen, S. C. & Tseng, T. Y. Parameter determination of support vector machine and feature selection using simulated annealing approach. Applied Soft Computing Journal 8, 1505–1512 (2008).

8. Mafarja, M. M. & Mirjalili, S. Hybrid Whale Optimization Algorithm with simulated annealing for feature selection. Neurocomputing 260, 302–312 (2017).

## References

1. Wolz, R. et al. Multi-method analysis of MRI images in early diagnostics of Alzheimer’s disease. PLoS ONE 6, 1–9 (2011).

2. Zhang, D., Wang, Y., Zhou, L., Yuan, H. & Shen, D. Multimodal classification of Alzheimer’s disease and mild cognitive impairment. NeuroImage 55, 856–867 (2011).

3. Liu, M., Zhang, D. & Shen, D. Ensemble sparse classification of Alzheimer’s disease. NeuroImage 60, 1106–1116 (2012).

4. Gray, K. R., Aljabar, P., Heckemann, R. A., Hammers, A. & Rueckert, D. Random forest-based similarity measures for multi-modal classification of Alzheimer’s disease. NeuroImage 65, 167–175 (2013).

5. Liu, X., Tosun, D., Weiner, M. W. & Schuff, N. Locally linear embedding (LLE) for MRI based Alzheimer’s disease classification. NeuroImage 83, 148–157 (2013).

6. Wee, C. Y., Yap, P. T. & Shen, D. Prediction of Alzheimer’s disease and mild cognitive impairment using cortical morphological patterns. Human Brain Mapping 34, 3411–3425 (2013).

7. Guerrero, R., Wolz, R., Rao, A. W. & Rueckert, D. Manifold population modeling as a neuro-imaging biomarker: Application to ADNI and ADNI-GO. NeuroImage 94, 275–286 (2014).

8. Payan, A. & Montana, G. Predicting Alzheimer’s disease a neuroimaging study with 3D convolutional neural networks. ICPRAM 2015 - 4th International Conference on Pattern Recognition Applications and Methods, Proceedings 2, 355–362 (2015).

9. Prasad, G., Joshi, S. H., Nir, T. M., Toga, A. W. & Thompson, P. M. Brain connectivity and novel network measures for Alzheimer’s disease classification. Neurobiology of Aging 36, S121–S131 (2015).

10. Suk, H. il, Lee, S. W. & Shen, D. Latent feature representation with stacked auto-encoder for AD/MCI diagnosis. Brain Structure and Function 220, 841–859 (2015).

11. Shakeri, M., Lombaert, H., Tripathi, S. & Kadoury, S. Deep spectral-based shape features for Alzheimer’s disease classification. Lecture Notes in Computer Science (including subseries Lecture Notes in Artificial Intelligence and Lecture Notes in Bioinformatics) 10126 LNCS, 15–24 (2016).

12. Aderghal, K., Benois-Pineau, J., Afdel, K. & Gwenaëlle, C. FuseMe: Classification of sMRI images by fusion of deep CNNs in 2D+e projections. ACM International Conference Proceeding Series Part F1301, (2017).

13. Aderghal, K., Boissenin, M., Benois-Pineau, J., Catheline, G. & Afdel, K. Classification of sMRI for AD Diagnosis with Convolutional Neuronal Networks: A Pilot 2-D+ ϵ Study on ADNI. in MultiMedia Modeling, Lecture Notes in Computer Science (eds. Amsaleg, L., Gu ðmundsson, G., Gurrin, C., Jónsson, B. & Satoh, S.) 690–701 (Springer International Publishing, 2017). doi:10.1007/978-3-319-51811-4_56.

14. Billones, C. D., Demetria, O. J. L. D., Hostallero, D. E. D. & Naval, P. C. DemNet: A Convolutional Neural Network for the detection of Alzheimer’s Disease and Mild Cognitive Impairment. IEEE Region 10 Annual International Conference, Proceedings/TENCON 3724–3727 (2017) doi:10.1109/TENCON.2016.7848755.

15. Guo, H., Zhang, F., Chen, J., Xu, Y. & Xiang, J. Machine learning classification combining multiple features of a hyper-network of fMRI data in Alzheimer’s disease. Frontiers in Neuroscience 11, 1–22 (2017).

16. Korolev, S., Safiullin, A., Belyaev, M. & Dodonova, Y. Residual and plain convolutional neural networks for 3D brain MRI classification. arXiv 835–838 (2017).

17. Wang, S., Shen, Y., Chen, W., Xiao, T. & Hu, J. Automatic Recognition of Mild Cognitive Impairment from MRI Images Using Expedited Convolutional Neural Networks. in Lecture Notes in Computer Science (including subseries Lecture Notes in Artificial Intelligence and Lecture Notes in Bioinformatics) vol. 10613 LNCS 373–380 (2017).

18. Li, F. & Liu, M. Alzheimer’s disease diagnosis based on multiple cluster dense convolutional networks. Computerized Medical Imaging and Graphics 70, 101–110 (2018).

19. Qiu, S. et al. Fusion of deep learning models of MRI scans, Mini–Mental State Examination, and logical memory test enhances diagnosis of mild cognitive impairment. Alzheimer’s and Dementia: Diagnosis, Assessment and Disease Monitoring 10, 737–749 (2018).

20. Senanayake, U., Sowmya, A. & Dawes, L. Deep fusion pipeline for mild cognitive impairment diagnosis. in 2018 IEEE 15th International Symposium on Biomedical Imaging (ISBI 2018) 1394–1997 (IEEE, 2018). doi:10.1109/ISBI.2018.8363832.

21. Altaf, T., Anwar, S. M., Gul, N., Majeed, M. N. & Majid, M. Multi-class Alzheimer’s disease classification using image and clinical features. Biomedical Signal Processing and Control 43, 64–74 (2018).

22. Forouzannezhad, P., Abbaspour, A., Cabrerizo, M. & Adjouadi, M. Early Diagnosis of Mild Cognitive Impairment Using Random Forest Feature Selection. in 2018 IEEE Biomedical Circuits and Systems Conference (BioCAS) vol. 53 1–4 (IEEE, 2018).

23. Hosseini-Asl, E., Gimel’farb, G. & El-Baz, A. Alzheimer’s disease diagnostics by a deeply supervised adaptable 3D convolutional network. Frontiers in Biosciences 23, 584–596 (2018).

24. Jie, B., Liu, M. & Shen, D. Integration of temporal and spatial properties of dynamic connectivity networks for automatic diagnosis of brain disease. Medical Image Analysis 47, 81–94 (2018).

25. Jie, B., Liu, M., Zhang, D. & Shen, D. Sub-Network Kernels for Measuring Similarity of Brain Connectivity Networks in Disease Diagnosis. IEEE Transactions on Image Processing 27, 2340–2353 (2018).

26. Raeper, R., Lisowska, A. & Rekik, I. Cooperative Correlational and Discriminative Ensemble Classifier Learning for Early Dementia Diagnosis Using Morphological Brain Multiplexes. IEEE Access 6, 43830–43839 (2018).

27. Basaia, S. et al. Automated classification of Alzheimer’s disease and mild cognitive impairment using a single MRI and deep neural networks. NeuroImage: Clinical 21, 101645 (2019).

28. Forouzannezhad, P., Abbaspour, A., Li, C., Cabrerizo, M. & Adjouadi, M. A Deep Neural Network Approach for Early Diagnosis of Mild Cognitive Impairment Using Multiple Features. Proceedings - 17th IEEE International Conference on Machine Learning and Applications, ICMLA 2018 1341–1346 (2019) doi:10.1109/ICMLA.2018.00218.

29. Wang, H. et al. Ensemble of 3D densely connected convolutional network for diagnosis of mild cognitive impairment and Alzheimer’s disease. Neurocomputing 333, 145–156 (2019).

30. Wee, C. Y. et al. Cortical graph neural network for AD and MCI diagnosis and transfer learning across populations. NeuroImage: Clinical 23, 101929 (2019).

31. Kam, T. E., Zhang, H., Jiao, Z. & Shen, Di. Deep Learning of Static and Dynamic Brain Functional Networks for Early MCI Detection. IEEE Transactions on Medical Imaging 39, 478–487 (2020).

32. Fang, C. et al. Gaussian discriminative component analysis for early detection of Alzheimer’s disease: A supervised dimensionality reduction algorithm. Journal of Neuroscience Methods 344, 108856 (2020).

33. Forouzannezhad, P. et al. A Gaussian-based model for early detection of mild cognitive impairment using multimodal neuroimaging. Journal of Neuroscience Methods 333, 108544 (2020).

34. Jiang, J., Kang, L., Huang, J. & Zhang, T. Deep learning based mild cognitive impairment diagnosis using structure MR images. Neuroscience Letters 730, 134971 (2020).

35. Kang, L., Jiang, J., Huang, J. & Zhang, T. Identifying Early Mild Cognitive Impairment by Multi-Modality MRI-Based Deep Learning. Frontiers in Aging Neuroscience 12, 1–10 (2020).

36. Yang, P. et al. Fused Sparse Network Learning for Longitudinal Analysis of Mild Cognitive Impairment. IEEE Transactions on Cybernetics 51, 233–246 (2021).

